# Calpain-2 regulates hypoxia/HIF-induced amoeboid reprogramming and metastasis

**DOI:** 10.1101/2020.01.06.892497

**Authors:** Veronika te Boekhorst, Liying Jiang, Marius Mählen, Maaike Meerlo, Gina Dunkel, Franziska C. Durst, Yanjun Yang, Herbert Levine, Boudewijn M. T. Burgering, Peter Friedl

## Abstract

Hypoxia, through hypoxia inducible factor (HIF), drives cancer cell invasion and metastatic progression in various cancer types, leading to poor prognosis. In epithelial cancer, hypoxia further induces the transition to amoeboid cancer cell dissemination, yet the molecular mechanisms, relevance for metastasis, and effective interventions to combat hypoxia-induced amoeboid reprogramming remain unclear. Here, we identify calpain-2 as key regulator and anti-metastasis target of hypoxia-induced transition from collective to amoeboid dissemination of breast and head and neck (HN) carcinoma cells. Hypoxia-induced amoeboid dissemination occurred through low ECM-adhesive, bleb-based amoeboid movement, which effectively invaded into 3D collagen with low-oxidative and -glycolytic energy metabolism, revealing an microenvironmentally-induced, energy-conserving dissemination route in epithelial cancers. Hypoxia-induced calpain-2 mediated amoeboid conversion by de-activating beta1 integrins, through enzymatic cleavage of the focal adhesion adaptor protein talin-1. Consequently, targeted downregulation of calpain-2 or pharmacological intervention restored talin-1 integrity, beta1 integrin engagement and reverted blebbing-amoeboid to elongated phenotypes under hypoxia. Calpain-2 activity was required for hypoxia-induced blebbing-amoeboid conversion in the orthotopic mouse dermis, and upregulated in invasive HN tumor xenografts in vivo, and attenuation of calpain activity prevented hypoxia-induced metastasis to the lungs. This identifies the calpain-2/talin-1/beta1 integrin axis as mechanosignaling program and promising intervention target of plasticity of cancer cell invasion and metastasis formation in epithelial cancers under hypoxia.

## Introduction

Cancer cell invasion initiates a multistep cascade to metastasis, which converts local neoplasia into a life-threatening systemic disease.^1–3^ Invading cancer cells migrate away from the primary tumor and penetrate blood and lymph vessels, followed by systemic spreading and metastatic colonization of distant organs.^1, 3^ For tissue invasion, cancer cells deploy a range of collective and individual cell migration strategies. Collective invasion of multicellular groups occurs when cells are held together by cell-cell adhesion, whereas single-cell migration lacks cell-cell cohesion and connectivity.^4^ Mesenchymal single-cell migration depends on effective integrin-mediated adhesion to the extracellular matrix (ECM), which supports spindle-like cell elongation and directs matrix metalloproteases (MMPs) for ECM remodeling and path generation.^5^ Amoeboid movement of roundish or ellipsoid cells engages only weak adhesion to the ECM, lacks ECM remodeling and, instead deploys kinetic deformation of the cell body for passage through 3D tissue.^6^ Besides the cell shape and the strength of ECM interactions, protrusion types differ between amoeboid migration modes. Amoeboid-moving leukocytes and cancer cells develop actin-rich pseudopodia and filopodia at the leading edge, which generate protrusive cell elongation and directional motion.^7, 8^ Alternatively, primordial germ cells and a subset of roundish-moving cancer cells extend bleb-like membrane protrusions which engage with the ECM and provide transient anchorage by their irregular surface topology and friction-based interaction.^7–11^ This repertoire of diverse physical and molecular cell-cell and cell-matrix interactions is present in normal cells as well as invasive cancer cells. While each migration mode operates according to distinct physicochemical principles, the relevance of each mode in the tumor microenvironment and during metastasis of cancer cells remains largely unclear.

Collective and single-cell migration programs are adaptive, allowing cells to switch between migration modes. Known conversions occur between collective and single-cell migration, including epithelial-to-mesenchymal or epithelial-to-amoeboid transition, as well as transitions between mesenchymal and amoeboid single-cell migration programs.^3^ Experimentally, migration mode switching can be induced by modulating mechanotransduction, e.g. inhibition of integrins or MMPs, or by increasing or decreasing actomyosin contractility.^12–15^ In tissues, adaptation of migration modes may also occur in response to physical and molecular stimuli, allowing metastatic tumor cells to optimize migration in complex and/or changing environments.^16^ The stimuli and mechanisms mediating epithelial-to-mesenchymal transition (EMT) have been investigated in detail. However, the pathophysiological requirements and intracellular regulators supporting the adaptive conversion to amoeboid migration are poorly understood.

Tumor hypoxia, a frequent microenvironmental stressor in solid tumors, induces invasion and metastasis in cancer cells^17–20^ and induces amoeboid migration in epithelial cancer cells.^21^ In response to severe hypoxia or pharmacological stabilization of hypoxia-inducible factor-1 (HIF-1), collectively invading cancer cells switch to amoeboid motion by largely EMT-independent mechanisms.^21^ Hypoxic and metabolically perturbed tumor regions surround necrotic zones in the tumor core but are also present in the tumor margins near the invasive front.^22^ After HIF-stabilization, invasive tumor cells retain sustained hypoxia-related transcriptional programs and metastatic ability.^22–26^ In response to hypoxic challenge, adaptive switching between migration modes may equip moving cancer cells with a repertoire of escape strategies, yet the molecular mechanisms mediating amoeboid conversion and their relevance for hypoxia-induced metastasis remain unclear.

We used invasive and metastatic epithelial breast cancer and head and neck squamous carcinoma cells (HN-SCC), which invade collectively into 3D fibrillar collagen in tumoroid culture upon normoxia and undergo rapid amoeboid conversion in response to hypoxia or pharmacological stabilization of HIF-1. We then mapped the induced amoeboid variants, pathways of migration mode switching, and their implications for metastatic organ colonization. Hypoxia/HIF stabilization induces conversion to blebbing-amoeboid movement by upregulating activity of the cysteine protease calpain-2, which cleaves the focal adhesion adapter talin-1, limits β1 integrin activity, and strongly enhances metastasis.

## Results

### Hypoxia induces blebbing amoeboid migration

To identify the amoeboid subtypes induced by hypoxia, 4T1 and UT-SCC38 cells that invade fibrillar collagen in 3D tumoroid culture by a predominantly collective pattern under normoxia (21% O_2_) were exposed to severe hypoxia (0.2% O_2_) or pharmacological HIF1-stabilization using DMOG (1 mM)^27^ (Fig. 1a, Supplementary Fig. 1a). After cells detached from the tumoroid, there were few spindle-shaped cells, and rounded migration via (i) actin-rich pseudopodal or (ii) blebbing protrusions toward the migration direction observed in control and, to a much greater extent, in hypoxic or DMOG-treated culture (Fig. 1b, arrowheads; Supplementary Fig. 1b, c). As a hybrid state, both blebs and pseudopodal or filopod-like protrusions coexisted in the same cell (Fig. 1b). The abundance of blebbing amoeboid migration among detached and individually migrating cells was 3- to 6-fold higher during hypoxia or HIF stabilization than under control conditions (Fig. 1c, Movie 1). The relative abundance of each migration mode differed between challenge conditions. In control culture, most detached single cells developed elongated, spindle-like shapes (E, 40%-60%), and smaller subsets developed amoeboid-like shapes with pseudopodal (P, 20%-30%) or bleb-based (B, 20%-30%) protrusions (Fig. 1d). Upon hypoxia or HIF stabilization, spindle-like (E, 10%-30%) and amoeboid pseudopodal (P, 10%-20%) subsets diminished, and the amoeboid blebbing mode more than doubled (B, 50%-80%) (Fig. 1d). This was confirmed by shape analysis of detached single-cell subsets, which showed the predominant abundance of roundish cells (EF <2) induced (150 % increase) by hypoxia, which is highly characteristic of amoeboid blebbing cells, which lack pointed pseudopodal protrusions (Supplementary Fig. 1 b,c). Migratory blebbing cells were proliferative (Supplementary Fig. 1d) and lacked signs of apoptosis, such as cleaved caspase-3, nuclear fragmentation (Supplementary Fig. 1e), cell collapse, and fragmentation (Movie 1). Thus, hypoxia-induced migratory blebbing phenotypes are viable. After HIF stabilization, all individual migration modes developed comparable median speeds (0.1 - 0.3 µm/min) in low- or mid-density collagen (1.7 or 2.5 mg/mL) (Supplementary Fig. 1f, g), and their speed and directional persistence was unperturbed when compared to rare individually moving cells in control culture (Supplementary Fig. 1f, h, i). In highest collagen density (4 mg/mL), all subtypes were strongly affected and slowed down, with blebbing amoeboid cells migrating with lowest velocity (Supplementary Fig. 1g). Thus, hypoxia induces a transition from collective to predominantly blebbing-amoeboid migration in epithelial cancer cells. This phenotype reaches optimal speed in low- to mid-density ECM, which is reminiscent of previously identified constitutive amoeboid movement in other experimental models of metastatic cancer dissemination.^10, 28^

**Figure 1.**
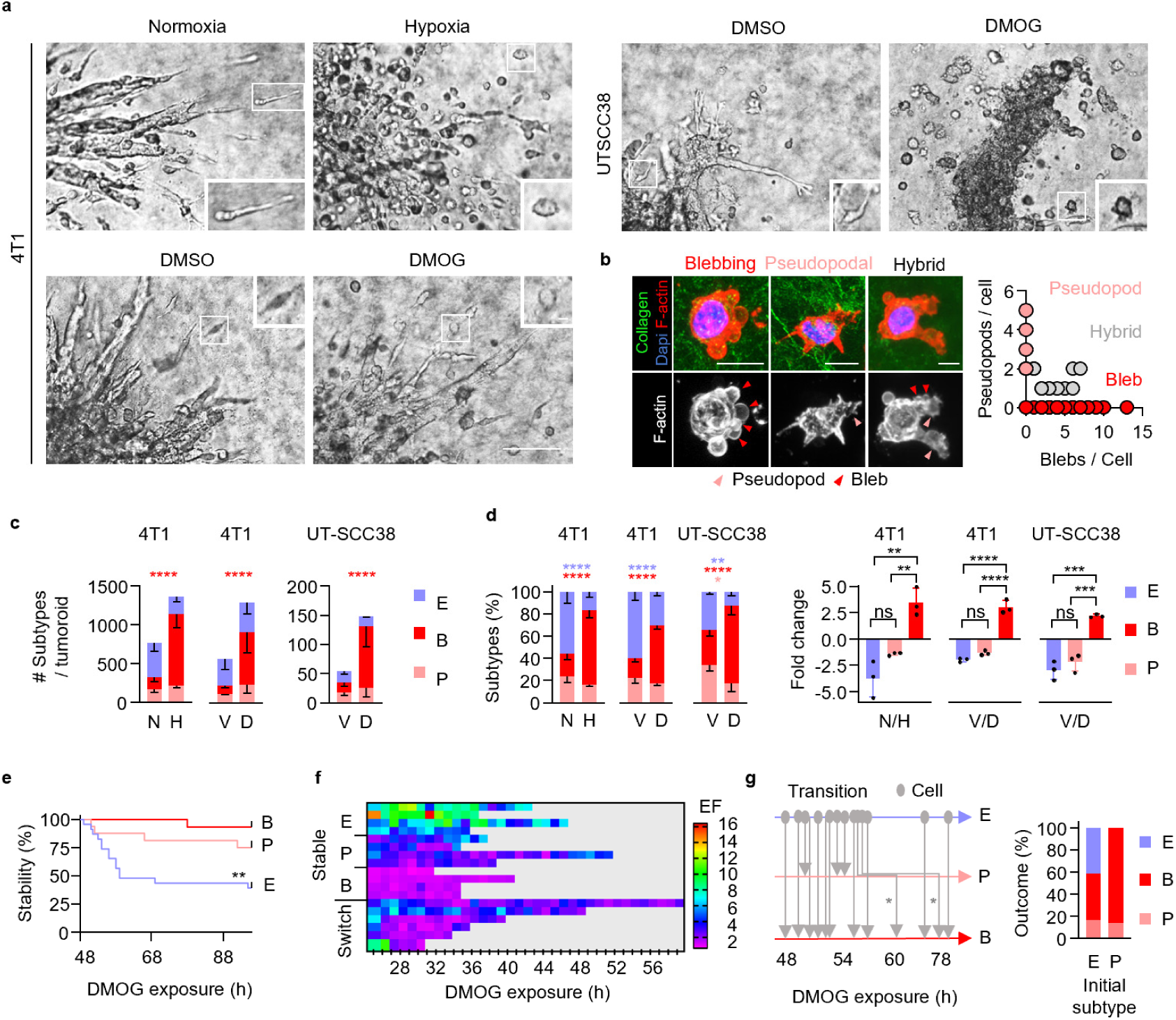
Hypoxia/HIF-induced amoeboid blebbing migration and cell viability. **a,** Brightfield micrographs of tumoroids invading into collagen at 72 h (4T1) and 96 h (UT-SCC38) under the conditions indicated. Insets, individually migrating cells. Scale bars, 100 μm (overviews), 10 μm (insets). **b,** Representative confocal images of UT-SCC38 cells 96 h after HIF-stabilization (left panels; inter-slice distance, 2 μm; scale bars, 10 μm). Protrusion type and number, including bleb-based, pseudopod-based and hybrid morphologies upon HIF stabilization (right panel; 44 cells, n=3). **c,** Morphology-based subtypes after 72 h (4T1) or 96 h (UT-SCC38). Data represent the experimental means ± s.d. of morphologic classifiers (5 tumoroids per condition each experiment, n=3). **** P<0.0001 (two-way ANOVA). **d,** Frequency of single-cell migration subtypes per tumoroid. Data are represented as in (**c**). **** P<0.0001, ** P<0.002, * P=0.015 (two-way ANOVA). Relative change of subtypes (right panel). **** P<0.0001, *** P<0.0008, ** P<0.008 (one-way ANOVA). **e,** Phenotypic stability of 4T1 migrating single cells after detachment from tumoroids. Migration modes were recorded by time-lapse microscopy (48 h observation period), represented as the time-dependent persistence or transition to another mode stable for > 3 h from 23 (E), 16 (P) and 13 (B) cells. ** P=0.002 (Log-rank test). **f,** Time-dependent elongation of individually moving 4T1 cells assigned to classified modes or switching behavior, represented as heat map. Elongation factor (EF), detected at 1-h intervals for the duration cells were in the migration field (typically 8-24h). Each row shows one cell monitored over time. **g,** Direct or step-wise (asterisks) transition of elongated to pseudopodal-round and to blebbing-round migration (left panel). Initial vs. resulting phenotypes of 4T1 single cells scored between 24 h and 48 h after HIF-stabilization (right panel; 38 cells). Abbreviations: E, elongated; P pseudopodal-rounded; B, blebbing-rounded; N, normoxia; H, hypoxia; V, vehicle control (DMSO); D, DMOG.

### Transitions and morpho-kinetic stability of blebbing amoeboid migration

Invasion plasticity induced by EMT^29, 30^ or adhesion-targeting antibody^15^ elicits collective to single-cell transition and results in long-lasting cell reprogramming (days). We thus tested the phenotypic stability of single-cell migration modes induced by hypoxia signaling. Within 48 to 72 hours after HIF stabilization, blebbing amoeboid migration in 4T1 cells was stable, without conversion to other phenotypes, whereas 20% to 50% of initially elongated or pseudopodal amoeboid cells were unstable and abandoned their mode (Fig. 1e). We used time-gated elongation analysis to detect the direction of mode switching of individual cells after HIF stabilization. While blebbing amoeboid cells retained a rounded shape (EF <2) over several days, pseudopodal (EF 1 - 4) or elongated, spindle-like cells (EF 3 to >10) frequently converted to round morphologies without notable reversion (Fig. 1f). As a consequence, after 3 days of HIF stabilization, collective, elongated, and pseudopodal amoeboid migration had transitioned to blebbing amoeboid migration in 4T1 cells (Fig. 1g, Movie 2) and UT-SCC38 cells (Fig. 1d, Supplementary Fig. 1j). These data reveal a microenvironmentally induced plasticity response where epithelial cancer cells transit from collective to blebbing-amoeboid migration, either directly or via transient elongated and pseudopodal states.

### Hypoxia and HIF stabilization reduce **β**1 integrin activity

Hypoxia-induced amoeboid conversion suggests decreased adhesion to the ECM substrate and/or increased actomyosin contractility through Rho/ROCK signaling, as shown in other models of cell detachment^15^ and amoeboid migration.^13, 31, 32^ To determine whether modulating adhesion, with or without modulating contractility, is sufficient to mediate rounded cell migration, we carried out mechanochemical simulations using a 2D phase-field model of cells moving across a 2D substrate (Supplementary Fig. 2a). This model has proven useful for determining the shape of keratocytes,^33^ the role of cytoskeletal contractility, and possible instabilities during cell motion.^34^ Cell elongation positively correlated with adhesion force and was decreased with reduced adhesion-dependent friction, particularly when contractility was increased (Fig. 2a). While actomyosin contractility is involved in collective, mesenchymal and amoeboid migration^4^, high actomyosin contractility along with low-adhesive interactions particularly accounts for amoeboid migration, where the cell body propulsively intercalates between ECM structures.^31^ We therefore hypothesized that hypoxia/HIF signaling may reduce cell adhesion to the ECM substrate and thereby cause rounding and amoeboid transition. Cell adhesion to type I collagen is predominantly mediated by integrins α1β1 and α2β1.^35^ Hypoxia or HIF stabilization both reduced the β1 integrin activation epitope in cell lysates of 4T1 cells (detected by mAb 9EG7) and UT-SCC38 cells (mAb TS2/16) in comparison to control cells, whereas total β1 integrin remained unchanged (Fig. 2b). To test whether decreased β1 integrin activation scales with weakened cell adhesion, weakly (detached) and firmly (attached) adherent cells were separated by applying shear force (mild washing with PBS) to hypoxic or normoxic 2D monolayer cultures (Fig. 2c, Supplementary Fig. 2b). HIF-stabilization downregulated active β1 integrin (by 50%) in detached cells, while adhesive cells retained high active β1 integrin levels (Fig. 2d, Supplementary Fig. 2c). Consistently, in hypoxic 3D invasion culture, β1 integrin activation epitope (9EG7) was decreased in individualized migrating cells, compared with normoxic culture (Fig. 2e, arrowheads). Hypoxia and HIF stabilization thus dampen β1 integrin activation in cell subsets that weakly adhere in 2D culture and undergo single-cell migration in 3D culture.

**Figure 2.**
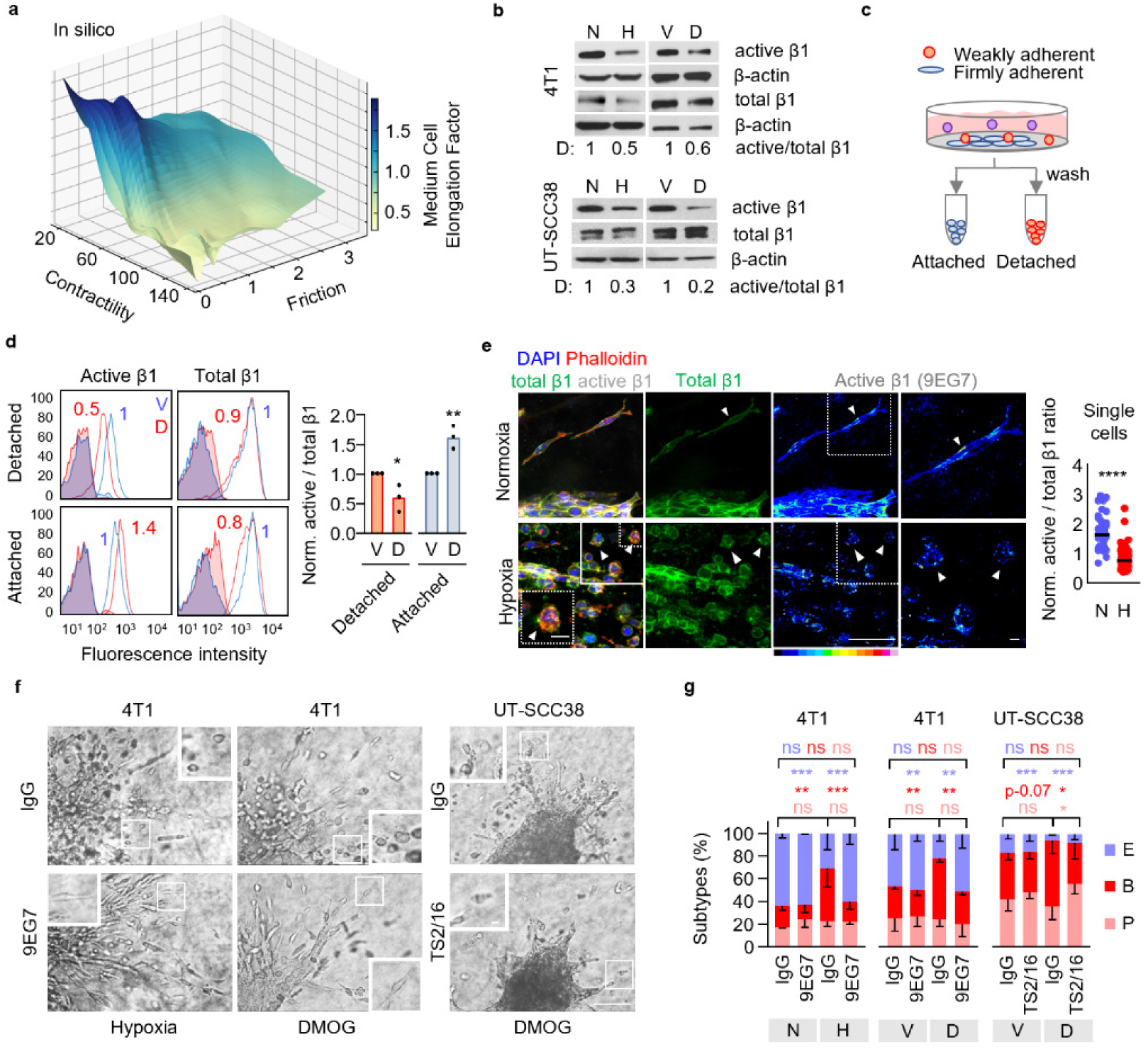
Reduced β1 integrin activity in amoeboid transition. **a,** In-silico modelling of cell elongation. Individual cell migration in dependence of friction and contractility using a two-dimensional phase field simulation. **b,** Active and total β1 integrin protein content (2D monolayer culture). D, densitometric analysis (representative Western blot, n=2-3). **c,** Schematic of cell isolation for harvesting attached (highly adhesive) and detached (weakly adhesive) cells in 2D culture. **d,** Active and total β1 integrin surface expression (MFI) in 4T1 subpopulations 48 h after treatment (left panel, representative flow cytometry histogram). Ratio of active/total β1 integrin surface expression normalized to vehicle control ratios (right panel). Columns show the median from independent experiments (data points). ** P=0.007, * P=0.03 (unpaired t-test, two-sided). **e,** Confocal micrographs of active (mAb 9EG7) and total β1 integrin (mAb CD29) expression of 4T1 tumoroids invading into 3D collagen (left panel) and the ratio of active/total β1 integrin per invading 4T1 single cell (right panel, 106 cells). Data show ratios of single cells; horizontal lines the median. Insets, single cell invasion phenotypes (arrowheads). Scale bars, 100 μm (overview), 10 µm (inset). **** P<0.0001 (Mann-Whitney test, two-sided). **f,** Brightfield micrographs of tumoroids after 72 h of 3D collagen invasion (left panel) and morphology-based single-cell subtypes (right panel) for indicated conditions in the presence or absence of β1 integrin-activating mAb 9EG7. Insets, morphologies of migrating single-cells. Data show the experimental means ± s.d. (5 tumoroids/condition each experiment, n=3). **** P<0.0001, *** P=0.0008, ** P<0.006, * P<0.03 (two-way ANOVA). Abbreviations: E, elongated; P pseudopodal-amoeboid; B, blebbing-amoeboid; N, normoxia; H, hypoxia; V, vehicle (DMSO); D, DMOG.

### Diminished **β**1 integrin activity is required for amoeboid conversion

Amoeboid migration occurs with weak or even absent integrin-mediated adhesion.^7, 31, 36^ We thus tested whether amoeboid migration in response to HIF stabilization was still supported by β1 integrins. The migration speed of detached single cells in the presence of adhesion-perturbing anti-β1 integrin mAb 4B4 was dose-dependently reduced, with 80% decline (from 0.1 µm/min to 0.02 µm/min) for maximum interference (Supplementary Fig. 2d). Notably, a moderate dose of mAb 4B4 mildly affected the speed of single UT-SCC38 cells, whereas it profoundly decelerated the migration speed of mesenchymal HT1080 fibrosarcoma cells (Supplementary Fig. 2d). This indicates that HIF-induced amoeboid migration retains moderate to weak β1 integrin-mediated adhesion. To test whether diminished β1 integrin function was required for the transition to amoeboid migration, β1 integrin was ectopically activated by mAb 9EG7 (4T1) or TS2/16 (UT-SCC38) added after HIF stabilization, when cells had detached from the tumoroid. Activation of β1 integrin decreased the frequency of round, bleb-rich phenotypes and enforced elongated morphologies with pseudopodal protrusions (Fig. 2f, g; Supplementary Fig. 2e, Movie 3). In contrast, integrin inhibition by mAb 4B4 induced round, blebbing migration in control culture, comparable to its occurrence after HIF stabilization (Supplementary Fig. 2f). These data indicate that hypoxia/HIF signaling induces bleb-based amoeboid migration by partially de-activating β1 integrin function.

### Elevated calpain-2 function mediates blebbing-amoeboid migration

Cell-intrinsic mechanisms that can limit integrin adhesion and weaken focal adhesion include (i) inhibition of intracellular phosphatases, which interrupts focal adhesion cycles^37^, and (ii) activation of the calcium-activated serine protease calpain-2, which cleaves the focal adhesion components talin, paxillin, and FAK.^38, 39^ Calpain-2 expression and activity are elevated throughout the metastatic cascade, including the primary tumor, lymph node metastases, and distant metastases of epithelial cancers such as colorectal cancer, squamous cell carcinoma, and breast cancer.^40–47^ We therefore explored the role of calpain as a putative regulator of integrin shutdown. Calpain-2 protein expression was mildly increased in hypoxic and DMOG-treated 2D culture compared with control conditions (Fig. 3a). In addition, the activity of intracellular calpain was increased by hypoxia and DMOG (up to 1.5 fold), and reverted by calpain-inhibitor PD150606 (Fig. 3b, c). Thus, calpain-2 expression and function are induced by hypoxia/HIF signaling.

**Figure 3.**
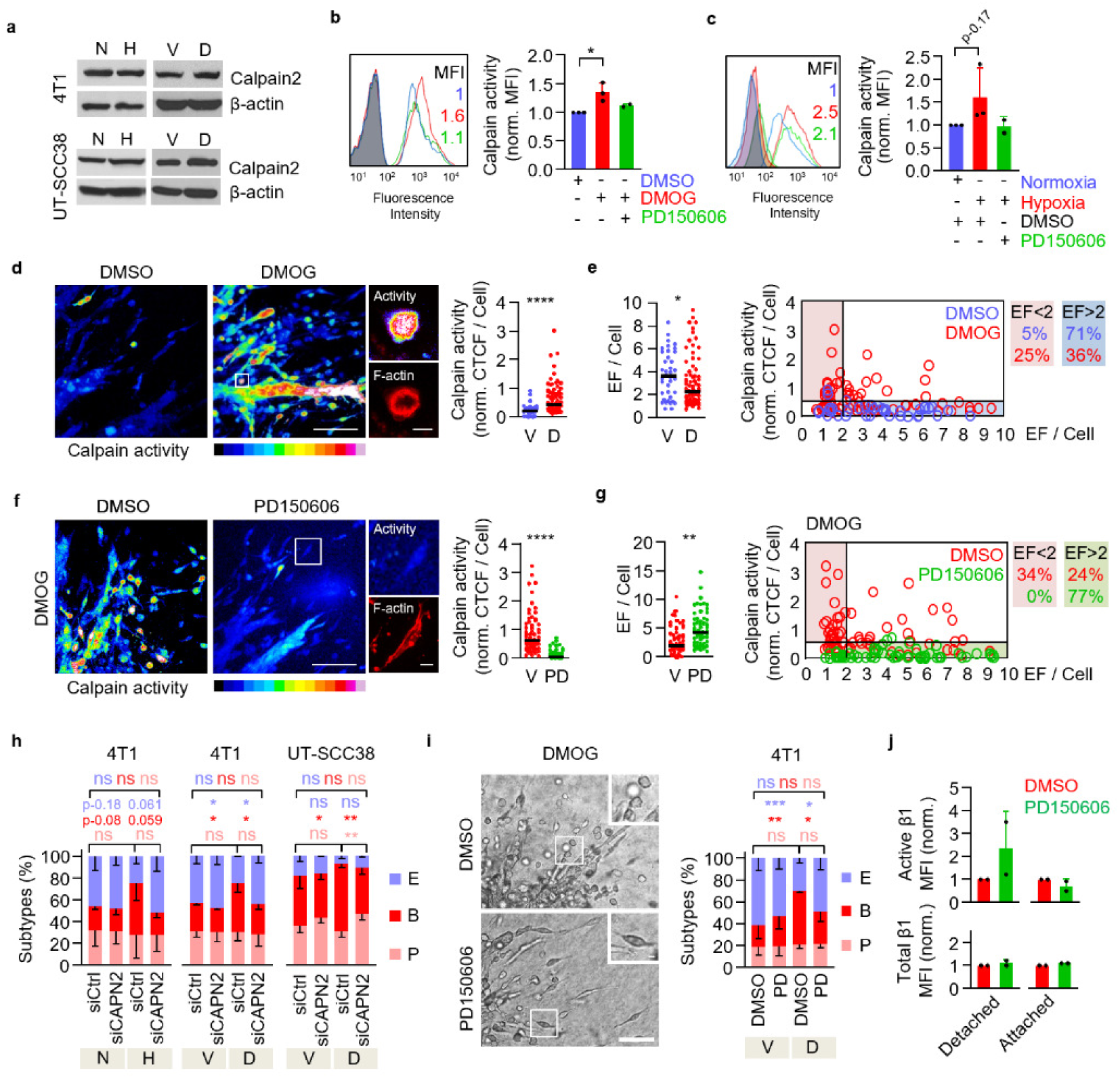
Calpain-2 dependent β1 integrin activation and amoeboid conversion after HIF stabilization. **a,** Calpain-2 expression after 72 h (4T1) and 96 h (UT-SCC38) in 2D culture under the indicated conditions. Representative Western blot (n=3). **b, c,** Calpain activity (cleaved CMAC intensity) detected by flow cytometry in 4T1 cells 48 h after HIF-stabilization with DMOG (**b**) or hypoxia (**c**), in the absence or presence of PD150606 (100 μM). Representative histograms with MFIs (left panels) and mean MFI ± s.d. normalized to controls (right panels; n=3). * P=0.02 (unpaired t-test, two-sided). **d,** Topography (left panel) and in situ cytometry (right panel) of calpain activity (cleaved CMAC intensity) in 4T1 single-cells detached from tumoroids in collagen 72 h after HIF-stabilization. Inset, amoeboid-blebbing cell; F-actin, phalloidin. CTCF, corrected total cell fluorescence from 118 cells pooled from n=2. **** P≤0.0001 (Mann-Whitney Test, two-sided). **e,** Elongation factor (EF) distribution (left panel) and correlation with calpain activity (right panel) of 4T1 single-cells invading in collagen. * P=0.028 (Mann-Whitney test, two-sided). **f,** Topography (left panel) and cytometry (right panel) of calpain activity of 4T1 tumoroids in collagen in the absence or presence of PD150606 (100 μM) after 72 h of HIF-stabilization. Data are represented as in (**d**). (72 h; 134 cells pooled from n=2). **** P≤0.0001 (Mann-Whitney Test, two-sided). **g,** EF distribution of 4T1 single cells (left panel) and correlation with calpain function (right panel) in the presence of PD150606 (100 μM) (right panel). Data show 134 cells from n=2. ** P=0.0051, * P=0.028 (Mann-Whitney test, two-sided). **h,** Migration morphologies of siCtrl- or siCAPN2-transfected single-cells 72 h (4T1) and 96 h (UT-SCC38) after treatment. Data represent the experimental means ± s.d. (5 tumoroids per condition each experiment, n=3). ** P<0.002, * P<0.05, ns P>0.95 (two-way ANOVA). **i,** Representative micrographs (left panel) and distribution of single-cell migration subtypes (right panel) of 4T1 tumoroids in collagen 72 h after HIF-stabilization with or without PD150606 (50 μM). Insets, single-cell migration subtypes. Data represent the experimental means ± s.d. (5 tumoroids per condition each experiment, n=3). *** P=0.0007, * P=0.05, ns P>0.31 (two-way ANOVA). **j,** Active and total β1 integrin surface expression in 4T1 cells after treatment with DMOG and PD150606 (100 μM). Data represent the mean MFI ± s.d. from n=2. Abbreviations: E, elongated; P, pseudopodal-amoeboid; B, blebbing-amoeboid; N, normoxia; H, hypoxia; V, vehicle (DMSO); D, DMOG; PD, PD150606. Horizontal lines (**d,e,f,g**), median. Scale bars (**d,f,i**), 100 μm (overviews), 10 μm (insets).

To link calpain activity to amoeboid conversion, we applied the fluorescent reporter t-BOC-L-Leucyl-L-Methionine amide (CMAC) to 3D tumoroid culture and quantified intracellular calpain activity in detached individual cells. Hypoxia and DMOG treatment caused calpain activation in both collective invasion strands and detached cells, whereas control cultures showed uniformly low activity (Fig. 3d, Supplementary Fig. 3a, b). Calpain activity elevation was high in HIF-induced rounded individual cells (EF <2) and low in elongated cells (EF >2) (Fig. 3e). This indicates that calpain activity is preferentially increased in amoeboid cells after conversion. Calpain inhibitor PD150606 diminished calpain activity to control level (Fig. 3f) and reverted cell rounding to elongated shapes despite ongoing HIF stabilization with DMOG (Fig. 3g).

To test whether the transition to blebbing amoeboid migration indeed depends on calpain-2 activity, we transiently downregulated calpain-2 by RNA interference and measured the effect on amoeboid conversion (Supplementary Fig. 3c, d). Calpain-2 downregulation prevented blebbing-amoeboid but induced elongated phenotypes in hypoxic or DMOG-treated culture (Fig. 3h; Supplementary Fig. 3e). Consistently, P150606 prevented HIF-induced amoeboid conversion in 3D tumoroid culture (Fig. 3i) and restored active β1 integrin surface levels in weakly adherent cells in 2D culture (Fig. 3j). Calpain-2 activation therefore dampens integrin activity and mediates HIF-induced amoeboid conversion.

### Calpain-2 activation mediates transition to amoeboid blebbing migration by talin-1 cleavage

We next addressed whether calpain-2 deactivates β1 integrins in HIF-induced moving cancer cells by its canonical function, i.e., cleaving focal adhesion complex proteins. Talin-1 is the central mechanosensitive adaptor protein in integrin-mediated focal contacts and an important calpain-2 target to induce focal adhesion turnover.^48, 49^ Hypoxia and HIF stabilization with DMOG both induced a proteolytic talin-1 fragment of 190kDa, which corresponds to the main calpain-2 mediated talin-1 cleavage product (Fig. 4a).^38, 39^ Talin-1 cleavage was reverted by 20% (hypoxia) and 70% (DMOG) after calpain inhibition with PD150606 (Fig. 4b) and after transient siRNA-mediated HIF/calpain-2–specific knockdown, where comparable levels to DMSO controls were reached (Fig. 4c, Supplementary Fig. 4a). Thus, hypoxia/HIF signaling induces proteolytic cleavage of talin-1 through calpain-2.

We finally tested whether the calpain-2/talin axis is required for transition to blebbing-amoeboid migration. GFP-TalinL432G, which resists calpain-mediated cleavage but retains wildtype talin function and stabilizes focal adhesions, was transiently expressed in 4T1 cells and the invasion modes were monitored in 3D tumoroid culture (Supplementary Fig. 4b). Ectopic expression of GFP-TalinL432G prevented cell rounding, abolished bleb formation, and induced spindle-shaped elongation despite HIF stabilization, whereas individually migrating GFP-transfected control cells remained rounded (Fig. 4d, e). GFP-TalinL432G focalized at the tips of cell extensions and along the cell body of elongated cells (Fig. 4f), consistent with focalized cell-matrix interactions.^48^ In summary, calpain-2 cleaves talin-1 to enable cell rounding and blebbing-amoeboid migration.

**Figure 4.**
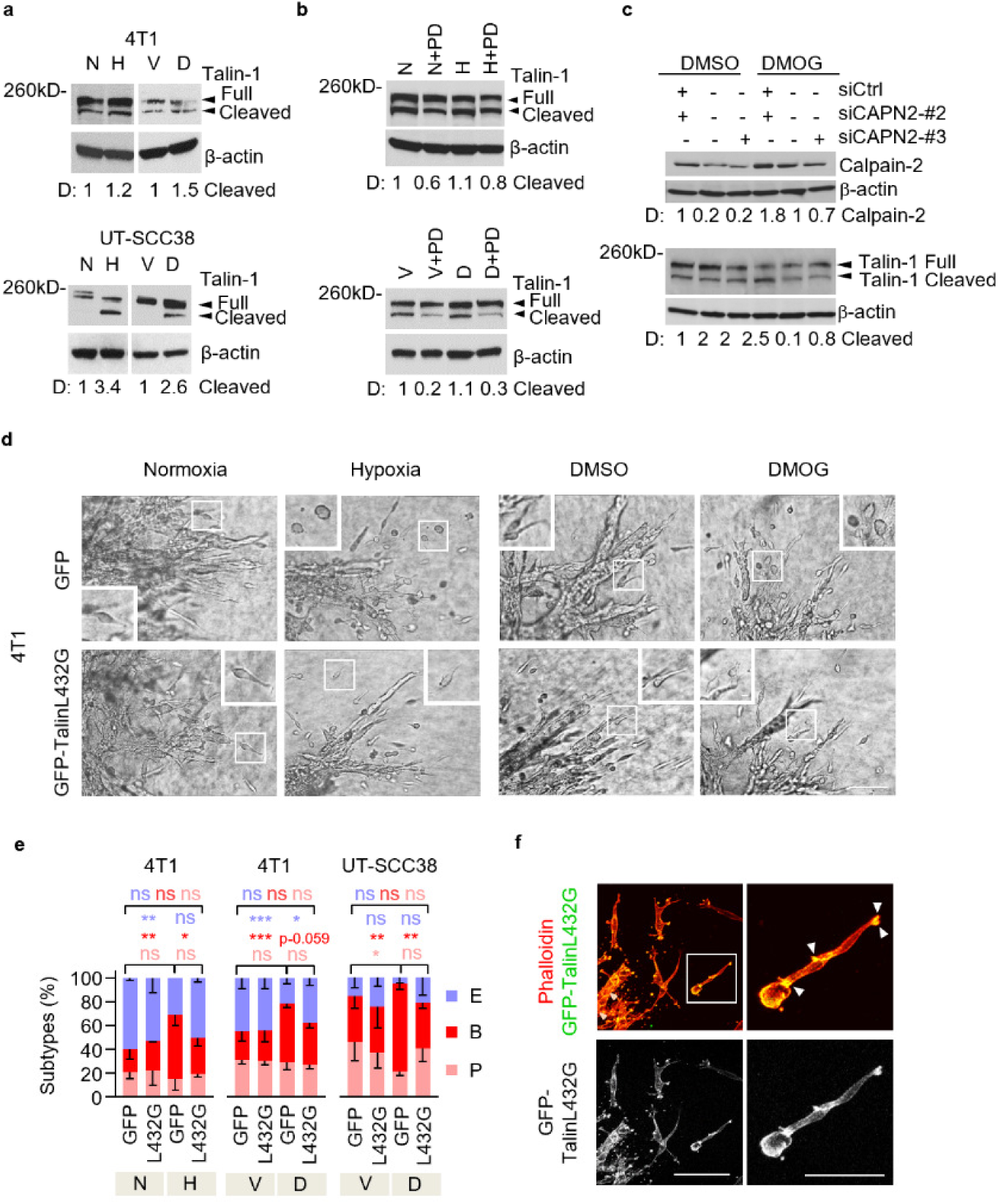
Calpain-2-induced talin-1 cleavage-dependent phenotypic conversion to amoeboid migration. **a,** Talin-1 protein expression, including full length (270 kDa) and cleavage fragment (190kDa), after 48-72 h (4T1) and 96 h (UT-SCC38) of hypoxic culture or HIF-stabilization with DMOG. Representative Western blots and densitometric analysis (D). n=2-3. **b,c,** Full-length and cleaved Talin-1 of 4T1 cells 48h after hypoxia/HIF-stabilization combined with calpain inhibition (PD150606, 100 μM) (**b**) and effects of transient siRNA (100nM)-mediated downregulation of calpain-2 (**c**). Representative Western blots and densitometric analysis (D). n=2-3. **d,** Representative brightfield micrographs of 4T1 tumoroids expressing GFP or calpain-uncleavable GFP-talin (T432L) invading into 3D collagen. Insets show single-cell migration phenotypes after 72 h of culture. Scale bar, 100 μm (overviews), 10μm (insets). **e,** Morphology of individualized cells expressing calpain-uncleavable talin (L432G) during hypoxia- or DMOG-treatment. Morphometric subtypes: E, elongated; P pseudopodal-amoeboid; B, blebbing-amoeboid. Data represent the experimental means ± s.d. (5 tumoroids per condition each experiment, n=3). *** P < 0.0009, ** P < 0.003 and * P < 0.04 (two-way ANOVA). **f,** Subcellular localization of uncleavable talin. Representative confocal micrographs of detached 4T1 cells expressing GFP-TalinL432G during migration in 3D collagen 72 h after transfection and simultaneous HIF-stabilization. Arrowheads, focalized clusters. Scale bar, 100 μm (overview), 50 μm (inset).

### Amoeboid conversion induced by HIF signaling coincides with a quiescent metabolic state

Cell migration is an energy consuming process.^50, 51^ However, hypoxia/HIF signaling typically occurs alongside nutrient deprivation and metabolic challenge, which may limit energy production required for cell migration.^52^ Because hypoxia/HIF challenge induced conversion to amoeboid movement with efficient single-cell speeds in low- to mid-density 3D fibrillar collagen (Supplementary Fig. 1 f, g), we hypothesized that amoeboid conversion may be associated with energy conservation to secure migration despite metabolic stress.

Glycolysis and mitochondrial respiration by oxidative phosphorylation represent key routes for cellular ATP production.^53^ To quantify metabolic adaptations in 3D culture, we generated collagen droplets containing one invading 4T1 tumoroid each and measured oxidative and glycolytic activity (Fig. 5a, Supplementary Fig. 5a). HIF stabilization strongly decreased oxygen consumption (OCR) in tumoroids (Fig. 5b), inducing a 50-70% reduction of basal and maximum mitochondrial respiration and mitochondrial ATP production (Fig. 5c). Likewise, HIF-stabilization decreased glycolytic activity measured by extracellular acidification (ECAR) in 3D tumoroid culture (Fig. 5d), while lactate release, the product of glycolysis, remained unchanged (Supplementary Fig. 5b). This suggests that HIF signaling in 3D tumoroid invasion culture decreases oxidative phosphorylation without compensatory increase of glycolysis (Fig. 5e), not unlike an energetically silent/resting state^54, 55^ and/or a switch to alternative energy sources (e.g. lipids).^56^

**Figure 5.**
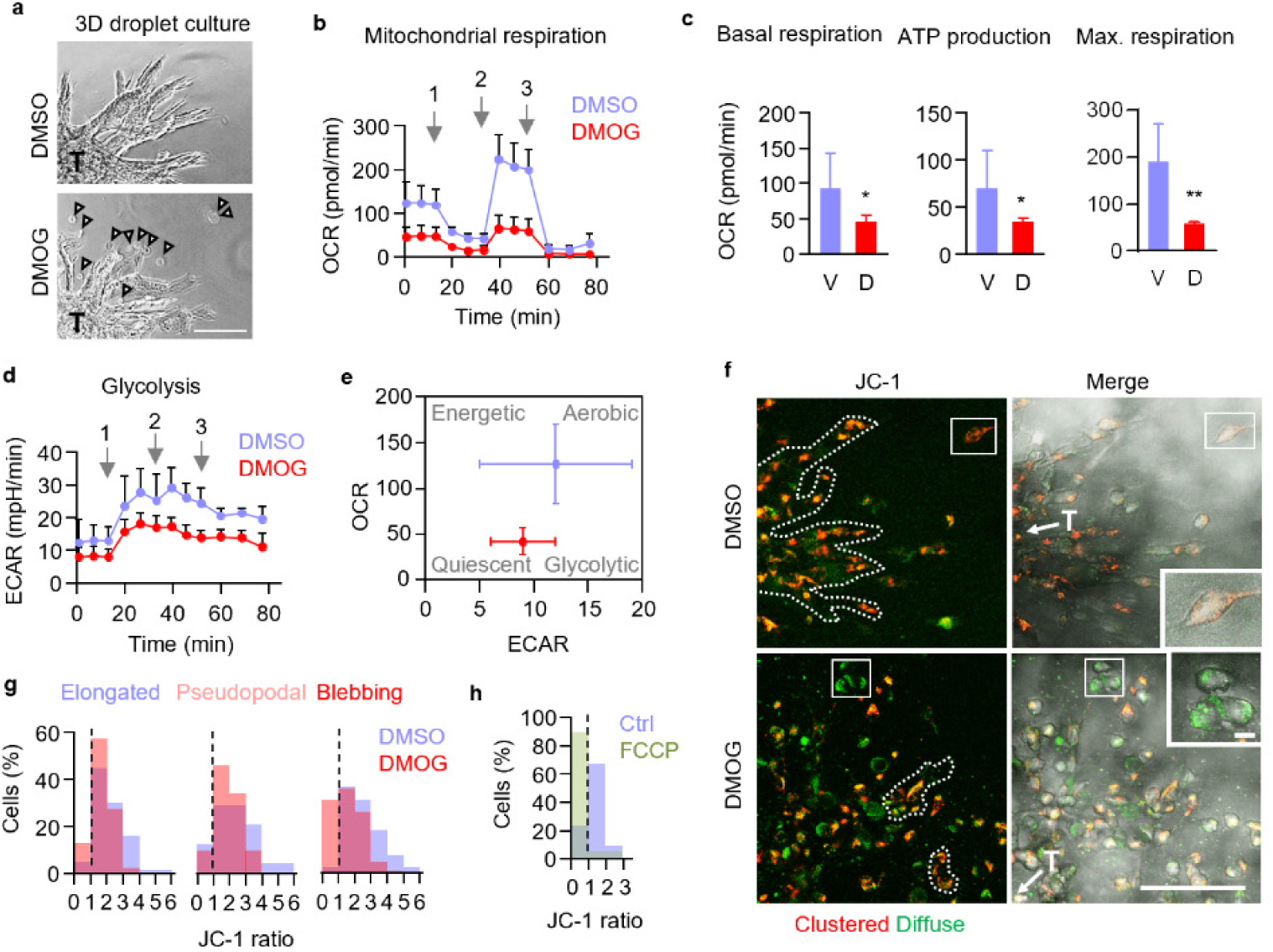
Metabolic adaptation associated with HIF-induced amoeboid reprogramming. **a,** Representative micrographs of 4T1 tumoroids (T) invading within collagen droplets used for Seahorse XF Mito Stress test. Arrowheads, detached invading single cells. Scale bar, 100 μm. **b,** Time-dependent oxygen consumption rate (OCR) of 4T1 tumoroids in collagen 48 h after HIF-stabilization. Arrows, time of addition of (1) oligomycin (2 μM), (2) FCCP (1 μM) and (3) rotenone and antimycin A (2 μM each). Data represent the means ± s.d. with 5-10 tumoroids per condition and experiment from n=3. **c,** Mitochondrial basal respiration, ATP production and maximum respiratory capacity of 4T1 tumoroids 48 h after HIF-stabilization measured by OCR. Data are represented as in (**b**). * P<0.03, ** P=0.005 (unpaired t-test, two-sided). **d,** Time-dependent extracellular acidification rate (ECAR) of 4T1 tumoroids 48 h after HIF-stabilization. Arrows and data are represented as in (**b**). **e,** Energy map of 4T1 tumoroids showing altered glycolytic (ECAR) and mitochondrial (OCR) activity after HIF-stabilization compared to control cultures. Date are compiled from (**b**) and (**d**). **f,** Representative confocal micrographs depicting mitochondrial activity detected by JC-1 (10 μg/ml) of 4T1 invasion into collagen 72 h after HIF-stabilization. Red fluorescence denotes high and green fluorescence low mitochondrial activity. T, tumoroid. Scale bars, 100 μm (overview), 10 μm (insets). **g,** Resulting JC-1 ratio (red/green) distribution within elongated (E), amoeboid-pseudopodal (P) and amoeboid-blebbing (B) 4T1 single-cell migration subsets from 103 (E), 74 (P) 137 (B) cells pooled from n=3. **h,** JC-1 ratio distribution of detached invading 4T1 single cells treated with a mitochondrial oxidative phosphorylation uncoupler FCCP (200nM) (63 cells, n=1).

To verify that mitochondrial activity decreased in migrating cells, we measured the mitochondrial membrane potential in 3D tumoroid culture using the ratiometric fluorophore tetraethylbenzimidazolylcarbocyanine iodide (JC-1) (Supplementary Fig. 5e). HIF stabilization induced a general reduction of the JC-1 ratio from 0.5-6 (control) to 0.5-4 (DMOG) (Fig. 5f, g), particularly in elongated and blebbing rounded-moving subsets (Supplementary Fig. 5c). Notably, a subset (30% of the total population) of amoeboid blebbing cells retained very low mitochondrial activity (JC-1 ratio <1), and this subset was absent in control cultures (Fig. 5g, dashed line; Supplementary Fig. 5c). A cell subset with similarly low mitochondrial activity was induced by oxidative phosphorylation uncoupling agent FCCP (Fig. 5h, dashed line). This suggests that blebbing amoeboid cancer cell subsets migrate with very low mitochondrial activity. HIF/calpain-induced amoeboid conversion thus enables an energy-conserving migration mode.

### Calpain activity, amoeboid conversion and relevance for metastasis *in vivo*

Calpain is activated in both early invasive primary tumors and metastatic lesions and further has been implicated in predicting poor prognosis in clinical epithelial cancers.^40–43, 47^ To explore whether calpain is activated in the invasion zone of epithelial tumors *in vivo*, we applied the calpain sensor CMAC to fresh-frozen sections of human HN-31 and Detroit562 HN-SCC after 6 to 8 weeks of orthotopic growth in nude mice. The invasion pattern of both tumors comprised multicellular nests and groups, typical for cross-sectioned collective invasion strands (Fig. 6a, Supplementary Fig. 6a),^57^ as well as adjacently positioned disseminated individual cells, which displayed notable rounded morphology (median EF 1.2 to 1.5) (Fig. 6a, arrowheads; Fig. 6b, Supplementary Fig. 6a). Compared to stromal regions, calpain activity was elevated in all tumor portions, including scattered multicellular nests and rounded tumor cells (Fig. 6c, Supplementary Fig. 6a). Thus, similar to collective strands and detached single cells in 3D culture, calpain activity is high in rounded, amoeboid-like cells in HN-SCC tumors.

**Figure 6.**
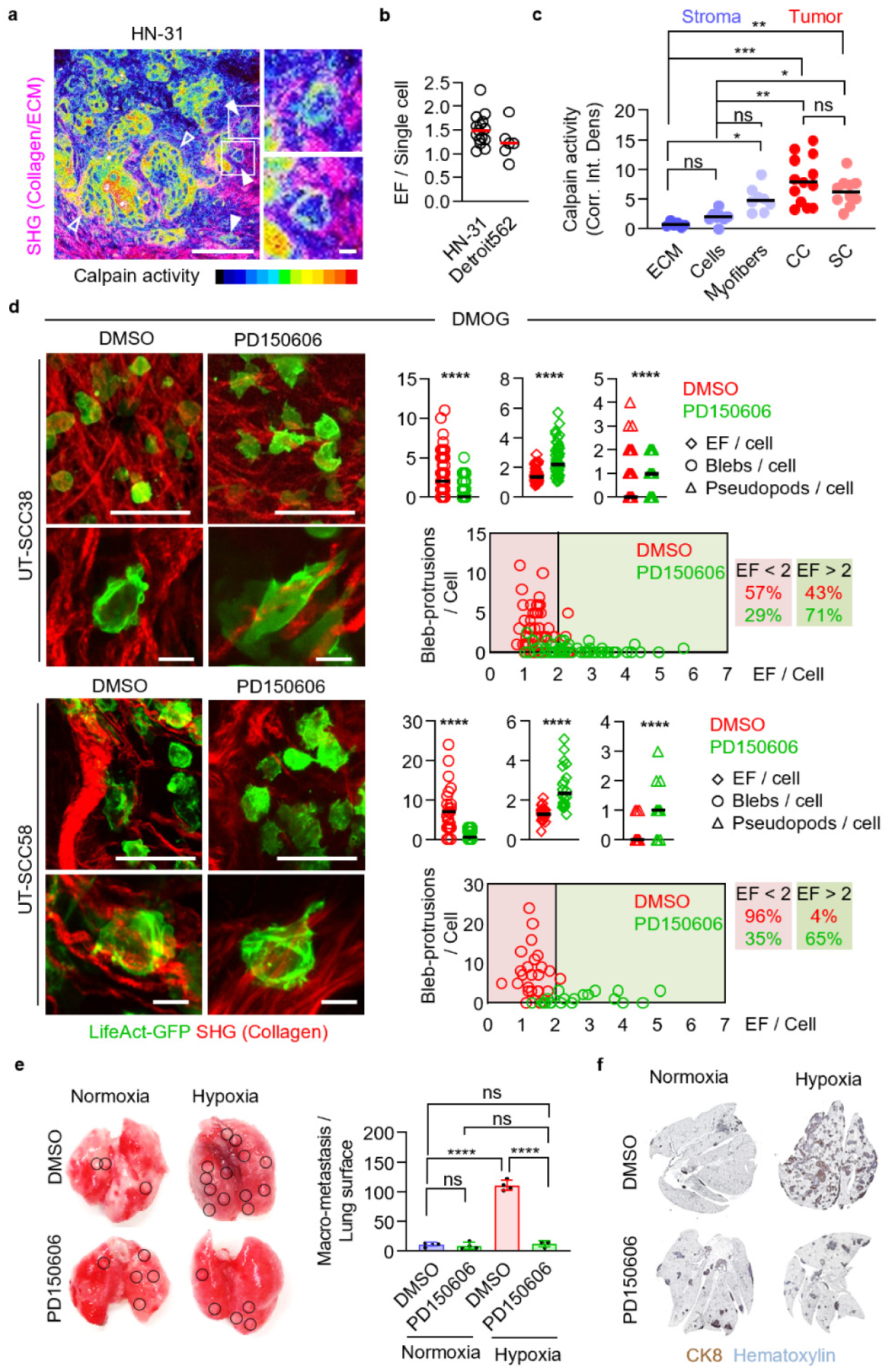
Calpain activity and function during tissue invasion and experimental metastasis in vivo. **a,** Calpain activity (cleaved CMAC intensity) in ex vivo orthotopic human HN-SCC tumor xenograft tissue. Insets and filled arrowheads, invading single cells with round morphology positive for cleaved CMAC. Open arrowheads, multicellular clusters detected by multiphoton microscopy. Scale bars, 100μm (overview), 10 μm (insets). **b,** Elongation factor (EF) distribution of invading single cells from orthotopically grown HN-SCC xenografts. Horizontal line, median (15 HN-31 from 4, 5 Detroit562 cells from 1 region, 1 tumor sample). **c,** Calpain activity (cleaved CMAC intensity) in tumor versus stromal compartments of human HN-31 HN-SCC tumor xenograft tissue sections. Data represent individual cells; horizontal lines, median. *** P=0.0001, ** P<0.003, * P<0.04 (Kruskal-Wallis test). **d,** Representative multi-photon images (left panels) and morphology and protrusion type analysis (right panels) of Lifeact-GFP expressing UT-SCC38 and UT-SCC58 single-cells migrating in the mouse dermis (SHG signal, collagen) after pre-treatment with DMOG combined with PD150606 (100 μM) or with solvent (DMSO) prior to intradermal injection (workflow depicted in Supplementary Fig. 6b). Data represent individual cells; horizontal lines, medians. (111 UT-SCC38, 45 UT-SCC58 cells, four tumor regions each). **** P<0.0001 (Mann-Whitney test, two-sided). Scale bars, 50 µm (overviews), 10 μm (insets). **e,** Macro-metastases (circles) on the lung surface of mice (left panel) and quantification (right panel) 14 days after tail-vein injection of 4T1 cells pretreated with normoxia and hypoxia with or without calpain inhibitor (PD150606, 100 μM; workflow depicted in Supplementary Fig. 6d). Columns show the mean ± s.d, with data points from individual lungs. **** P<0.0001 (one-way ANOVA). **f,** Representative micrographs from mid-sections of whole lungs bearing 4T1 metastasis, identified by cytokeratin-8 staining.

To directly test whether calpain controls cell shape *in vivo*, locally invasive but non-metastatic UT-SCC38 and invasive and metastatic UT-SCC58 cells were pretreated with HIF stabilizer DMOG, orthotopically implanted into the collagen-rich mouse dermis, and monitored by intravital microscopy (Supplementary Fig. 6b). DMOG-pretreated cells developed round shapes with multiple blebs in contact with collagen fibrils and bundles (Fig. 6d, Supplementary Fig. 6c), and some cells intercalated between collagen structures (Supplementary Fig. 6c). Coexposure with DMOG and calpain inhibitor PD150606 diminished bleb-like protrusions and favored elongated morphologies with pseudopod-like protrusions in contact with fibrillar collagen (Fig. 6d, Supplementary Fig. 6c). These data indicate the presence of calpain-dependent blebbing amoeboid phenotypes in dermis tissue *in vivo*.

Hypoxia is a strong inducer of metastatic activity,^58^ and calpain-2 activation predicts disease progression and poor prognosis in epithelial cancer.^47^ To investigate whether hypoxia-induced metastasis depends on calpain, 4T1 cells were maintained in hypoxic or normoxic two-day culture *in vitro* in the absence or presence of calpain inhibitor PD150606, injected into the tail vein of nude mice, and analyzed for lung metastasis after 14 days (Supplementary Fig. 6d). Hypoxia enhanced lung metastasis profoundly, and this effect was reverted by low-dose calpain inhibition to baseline levels obtained by normoxic control culture (Fig. 6e, f). Exposure to PD150606 did not compromise cell viability or proliferation rates during pretreatment (Supplementary Fig. 6e, f) nor the colony-forming capacity *in vitro* (Supplementary Fig. 6g). Thus, calpain activity is a key regulator of hypoxia-induced metastatic organ colonization and outgrowth.

## Discussion

Tumor hypoxia *in vivo* occurs before and during invasion at the primary tumor site, and likely in metastatic lesions.^22, 23^ Combining *in vitro, in vivo,* and *in silico* analysis of epithelial cancer cell migration, we here show that hypoxia/HIF signaling provides a microenvironmental trigger for rapid plasticity of mechanocoupling and adaptation of invasion, i.e., the transition from collective or mesenchymal to blebbing amoeboid migration. Amoeboid conversion enables a low-adhesive and energy-conserving migration strategy, which secures cell displacement despite hypoxic challenge. By activation of calpain-2 and cleavage of talin-1, HIF signaling dampens β1 integrin activity and thereby induces blebbing amoeboid migration and metastatic lung colonization as calpain-dependent processes. Consequently, pharmacological inhibition of calpain activity represents a promising strategy to limit or prevent HIF-induced amoeboid conversion and metastasis.

We here show that HIF-induced blebbing-amoeboid conversion is distinct from, and should not be confused with, morphologically similar apoptotic blebbing, based on: (i) intact cell viability over several days, (ii) ongoing occasional cell proliferation, (iii) asymmetry of blebs to the leading cell edge providing polarized interactions with collagen fibers in the direction of migration (compare Movie 2), (iv) induction of the calpain/talin axis but not of caspase activity, and (v) the ability of blebbing cells to translocate up to hundreds of micrometers per day. Amoeboid blebbing of epithelial cancer cells is thus associated with cell survival and migration. Therefore, our results support the concept that blebbing amoeboid migration is a fundamental and evolutionary conserved cell migration program detected in (patho)physiological contexts. These include *Dictyostelium discoideum* amoebae under nutrient starvation,^59^ migrating primordial germ cells forming the gonade in zebrafish embryos,^60^ melanoma cells,^61–63^ and as shown here, epithelial cancer cells in response to hypoxic perturbation.

A range of experimental *in vitro* interventions have helped to establish the concept that mesenchymally and collectively invading cancer cells can undergo amoeboid conversion, including inhibition of MMPs, extreme confinement, targeted inhibition of integrins or deprivation of ECM ligands, and augmenting Rho-mediated actomyosin contractility.^11, 15, 31, 64, 65^ We here show that hypoxia signaling represents a physiologically relevant upstream stimulus that limits integrin function and induces blebbing amoeboid dissemination in 3D tissues. HIF signaling is induced by a range of microenvironmental triggers, including hypoxia and acidosis,^66^ and/or growth factor signaling and cell transformation through PI3K/Akt/mTOR and MEK/ERK pathways.^67^ Thus, beyond hypoxia, HIF signaling may trigger amoeboid escape in response to various cell-autonomous and microenvironmental stimuli.

As an underlying mechanochemical program, the calpain-2 / talin-1 axis was sufficient to downregulate integrin activity in epithelial cancer and induce low-adhesive amoeboid migration. Low-adhesive migration is a well-established cell-displacement strategy: leukocytes retain constitutive, largely integrin-independent migration,^7^ and rounded cancer cells migrate despite targeted adhesion perturbation.^12, 64^ Calpain-2 regulates focal adhesion turnover in the cell rear, e.g., in adhesive fibroblasts, where it locally weakens adhesion bonds and facilitates uropod sliding without affecting overall cell adhesion and elongation.^38, 40^ By contrast, hypoxia-induced calpain-2 activation seems to downregulate integrin function throughout the cell, affecting both leading and trailing cell poles and thereby enabling whole-cell deadhesion and amoeboid conversion. Thus, calpain-2 serves as regulator of global cell adhesion and inducer of migration plasticity, enabling migration mode switching as a cell-intrinsic process. Beyond regulating the integrin adhesome, calpain-2 cleaves other proteins involved in cell migration, including cadherins and β-catenin,^38^ hence weakening cell-cell junctions and favoring individualization.

We further identified amoeboid-blebbing motion as an energy conserving migration strategy. Actomyosin treadmilling and focal adhesion formation, signaling and turnover are ATP consuming processes important in mesenchymal migration.^68^ Amoeboid movement retains actomyosin contractility and cortical actin dynamics, while integrin and protease systems are dispensable,^11^ thus potentially minimizing the energy demands of migration by modular activation/deactivation of non-obligatory activity. By minimizing energy demands, HIF/calpain-2-induced blebbing amoeboid migration may thus secure cell escape from metabolically perturbed tissue sites. This supports the concept that cancer cells adapt their migration paths^51^ and, as shown here, their migration mode according to energetic requirements.

Our data associate HIF/calpain-2–induced dissemination strategies with systemic dissemination and metastasis. However, calpain activity levels in tumor versus stromal sub-compartments remain unexplored. By applying the CMAC calpain activity probe for live-cell and ex vivo histochemistry, we found calpain activity upregulated during collective and single-cell invasion, i.e., in disseminated rounded cancer cell subsets. Detecting calpain activity across migration modes may thus identify intratumor niches of particular invasiveness and dissemination ability, as well as amoeboid conversion.

Besides transcriptional regulation of CAPN-2 as an HIF target gene, ^69–71^ hypoxia/HIF may trigger calpain activity through multiple routes, including regulation of the small regulatory subunits S1 or S2 of calpain or its endogenous inhibitor calpastatin, intracellular calcium transients, or protein phosphorylation.^38^ While the upstream mechanisms that regulate calpain activity in response HIF signaling remain to be shown, our data suggest that the HIF-induced calpain response is durable and supports hypermetastatic ability beyond the period of hypoxic challenge. Under continuous hypoxia, blebbing amoeboid migration remained stable over several days, without reversion to other migration modes. HIF stabilization, as pretreatment regime, secured blebbing invasion in the normoxic dermis over several hours and mediated metastatic lung colonization over several days, despite discontinuation of hypoxia/HIF trigger. Consistently, drug-based inhibition of calpain-2 prevented HIF-induced amoeboid conversion and metastasis. Thus, calpain-2 is a promising target for limiting HIF-promoted metastatic invasion and for prevention of secondary metastasis in high-risk patients and patients with a high load of circulating tumor cells.

## Online Methods

### Cell culture

Murine 4T1 breast carcinoma cells (CRL-2539, ATCC), a model for local invasion and distant metastasis,^72^ were cultured in RPMI (RPMI1640; Sigma Aldrich) containing 10% fetal bovine serum (FBS; Sigma Aldrich), penicillin (100 U/mL), streptomycin (100 µg/mL; PAA), and sodium pyruvate (1 mM; Thermo Fisher Scientific). Human UT-SCC38 (fully epithelial, locally invasive, non-metastatic) and UT-SCC58 (locally invasive, metastatic) head and neck squamous carcinoma cells (HNSCC),^73^ as well as UT-SCC38 Lifeact-GFP and UT-SCC58 Lifeact-GFP cells, were cultured in Dulbecco’s modified Eagle medium (DMEM; Sigma Aldrich), supplemented with 10% FBS (Sigma Aldrich), penicillin (100 U/mL), streptomycin (100 µg/mL; PAA), and sodium pyruvate (1 mM; Gibco).

Human Detroit 562 (ATCC) and HN-31 (provided by Dr. John Ensley, Wane State University, Detroit, MI) HNSCC cell lines were cultured in DMEM (Gibco) supplemented with 10% FCS (Sigma Aldrich). Stable Detroit 562 cell lines expressing control vector for short hairpin RNAs (shRNAs) were established as described previously.^74^ The tumor cell identity was verified by short tandem repeat (STR) DNA profiling (IDEXX BioResearch) and no interspecies contamination was detected. Cells were routinely tested negative for mycoplasma contamination (Myco Alert, Lonza).

### Reagents

The following antibodies and dyes were used: rabbit polyclonal anti-HIF1α (1:1000 WB, Novus Biologicals, 100-479), rabbit polyclonal anti–calpain-2 large subunit (1:1000 WB, Cell Signaling, 2539), chicken polyclonal anti-β-actin (1:1000 WB, Abcam, 13822), rabbit polyclonal anti-β-actin (1:2000 non-reducing WB in 4T1, Cell Signaling, 4967), rabbit polyclonal anti–talin-1 (clone 8d4; 1:200 WB, Sigma-Aldrich, T3287), mouse monoclonal anti-human CD29 (activated β1-integrin, clone HUTS-4; 1:500 WB, MerckMillipore, 2079Z), mouse monoclonal anti-human CD29 (clone 4B4; 1:1000 WB, 0.5-10 μg/ml functional studies, Beckman Coulter, 6603113), rat monoclonal anti-mouse CD29 (primed β1-integrin, clone 9EG7; 1:500 WB, 1:25 FC, 1:50 IF, BD Biosciences, 553715), rabbit monoclonal anti-mouse CD29 (clone EP1041Y; 1:1000 WB, Millipore, 04-1109), rat IgG2a isotype control (Clone R5-95; 1:25 FC, BD Biosciences, 553927), FITC-conjugated Armenian hamster anti-CD29 (clone HMβ1-1; 1:100 IF, 1:50 FC, Biolegend, 102206), FITC-conjugated Armenian hamster IgG (CloneHTK888; 1:20 FC, Biolegend, 400906), rabbit anti-cleaved caspase-3 (1:200 IF, Cell Signaling, 9664), mouse anti-human CD29 (clone TS2/16; 20 μg/ml, BioLegend, 303010), rabbit-on-rodent HRP-polymer (IHC, Biocare Medical, RMR622), rabbit monoclonal anti-cytokeratin 8 (clone EP1628Y; 1:250 IHC, Abcam, 53280), secondary goat anti-rabbit/mouse/chicken antibodies conjugated to horseradish peroxidase (1:10,000 WB, Jackson, 211-032-171/115-0450174/ 103-035-155), secondary goat anti-rat IgG AF488 (1:200 FC, Thermo Fisher Scientific, 11006), secondary goat anti-rat IgG AF647 (1:200 IF, Thermo Fisher Scientific, A21247), secondary goat anti-rabbit IgG AF633 (1:200 IF, Thermo Fisher Scientific, A21071), phalloidin AF488/546/568/633 (1:100 IF, Thermo Fisher Scientific, A12379/A22283/A12380/A22284), DAPI (1:500 IF, Thermo Fisher Scientific, D21490).

The following inhibitors and reagents were used: PD150606 (PD; Cayman Chemicals), etoposide (Sigma Aldrich), oligomycin (Bio-Connect), carbonyl cyanide-4-(trifluoromethoxy)phenylhydrazone (FCCP, Sigma Aldrich), Rotenone (Sigma Aldrich), Antimycin a (Sigma Aldrich), Dimethyl oxalylglycine (DMOG; Cayman Chemicals), dimethyl sulfoxide (DMSO; Sigma Aldrich), tetraethylbenzimidazolylcarbocyanine iodide (JC-1; Sigma Aldrich), t-BOC-L-leucyl-L-methionine amide (CMAC; Thermo Fisher Scientific), Hanks balanced salt solution (HBSS; with magnesium and calcium, Gibco) trypsin-EDTA (Sigma), methyl cellulose (Sigma Aldrich), collagen I rat-tail (Corning), bovine collagen solution (Advanced Biomatrix), LIVE/DEAD fixable stain (Thermo Fisher Scientific), para-formaldehyde solution (PFA; Affymetrix), L-glutamine (Sigma Aldrich), pyruvate (for metabolic flux analyses, Sigma Aldrich; for cell culture, Gibco), glucose (Sigma Aldrich), MISSION siRNA transfection reagent (Sigma Aldrich), Lipofectamine 2000 (Thermo Fisher Scientific), OptiMEM (Thermo Fisher Scientific), Rodent Decloaker (Biocare Medical), TBS Wash Buffer 1X (Biocare Medical), sodium oxamate (Sigma Aldrich), protease inhibitor cocktail (Roche), Bradford (Biorad), Novex 4-20% Tris-Glycine Mini Gels (Thermo Fisher Scientific), SQ-PVDF membranes (Thermo Fisher Scientific), SuperSignal West Pico Chemiluminescent Substrate (Thermo Fisher Scientific), CL-Xposure films (Thermo Fisher Scientific).

The following CAPN2-targeting siRNAs and plasmids were used: MISSION siRNA targeting mouse CAPN2 (NM_009794; siRNA ID-#1: SASI_Mm01_00183464, sequence start 1243, siRNA ID-#2: SASI_Mm01_00183465, sequence start 1820, siRNA ID-#3: SASI_Mm01_00183467, sequence start 2132; Sigma Aldrich), MISSION siRNA targeting human CAPN2 (NM_001748, siRNA ID-#1: SASI_Hs01_00059356, Sequence start 2234, siRNA ID-#2: SASI_Hs01_00059357, Sequence start 1053, siRNA ID-#3: SASI_Hs01_00059364, Sequence start 810, Sigma Aldrich), MISSION siRNA universal negative control (SIC001; Sigma Aldrich), plasmid DNA construct GFP-TalinL432G (26725; Addgene), pEGFP N1 (vector control; Clontech Laboratories).

### Hypoxia culture and DMOG treatment

For cell function studies, cells were cultured as 2D monolayers or tumoroids in 3D collagen lattices in normoxia (21% O2) or hypoxia (0.2% O_2_, CB53 incubator, Binder GmbH) for 72 hours (4T1) or 96 hours (UT-SCC38),^21^ unless indicated otherwise. For pharmacological activation of HIF in culture, cells were treated with DMSO (0.1, solvent control) or the prolyl-hydroxylase inhibitor DMOG (1 mM) for 72 hours (4T1) or 96 hours (UT-SCC38).

### Cell isolation from 2D culture

Subconfluent monolayers were grown under normoxic and hypoxic conditions or DMSO and DMOG for 72 hours (4T1) or 96 hours (UT-SCC38), unless otherwise stated. For separation of low- and high-adhesive cell populations, the supernatant medium was gently removed and low-adhesive cells were harvested by manual horizontal circular washing with warm PBS (100 cycles, 37 °C). Cells that remained plate-attached after washing (i.e., the highly adhesive population) were detached using trypsin-EDTA (2mM, 37 °C), and the low- and high-adhesive subsets were used separately or pooled for further analyses (cell viability, flow cytometry, Western blotting).

### Cell viability and proliferation analysis from 2D culture (treatment workflow underlying experimental metastasis assay)

4T1 cells were cultured under normoxia or hypoxia in the presence of calpain inhibitor PD150606 (100 μM) or solvent (DMSO) for 48 hours. DMSO and PD150606 were refreshed after every 20 hours. Cell viability was detected by Trypan Blue exclusion. The proliferation ratio reflects the total cell number harvested at the end-point divided by the number of cells seeded.

### Colony formation assay

A chilled well-plate was coated with a smooth bottom layer of reconstituted basement membrane (rBM) (matrigel; BD Bioscience) by centrifugation with slow acceleration (4 °C, 300 x g, 10 min) and solidification (37 °C, 30 min). 4T1 cells were cultured under normoxia or hypoxia in the presence of calpain inhibitor PD150606 (100 μM) or solvent (DMSO) for 2 days (refreshed every 20 hours and 2 hours before harvesting). Low- and high-adhesive cells were harvested (by PBS rinse and trypsin-EDTA (2 mM), respectively), pooled, resuspended in medium containing 5% rBM solution and added onto polymerized rBM for 5-day culture (37 °C, 5% CO_2_), with medium refreshment at day 3. Colonies were fixed (2 % pre-warmed PFA, 5 min), followed by incubation in 4 % PFA (10 min) and staining with DAPI. Colonies were imaged by epifluorescence and brightfield microscopy (Olympus IX-71, EM-CCD camera). Z-stacks were displayed as composites of maximum projections.

### Immunoblotting

Western blot analysis of protein expression was performed using subconfluent 2D cell monolayers. Cells were washed twice with ice-cold PBS, followed by protein extraction with lysis buffer (10 mM Tris, pH 8.0; 1 mM EDTA, pH 8.0; 150 mM NaCl; 0.1% NP-40), supplemented with a protease inhibitor cocktail. For detection of active β1 integrin epitope, cells were lysed under non-reducing conditions. Protein concentration was determined by the Bradford assay. Lysates were loaded on Novex 4%-to- 20% Tris-Glycine Mini Gels and transferred by electroblotting (120 V, 1 hour) in transfer buffer (28 mM Tris-HCL; 39 mM Glycine; 20% methanol v/v) onto SQ-PVDF membranes. Immunodetection was performed using primary antibody dilutions (1/100 – 1/1000) and HRP conjugated secondary antibodies (1/2000 – 1/10000 dilutions) in combination with the SuperSignal West Pico Chemiluminescent Substrate and CL-Xposure films. Densitometry of bands was performed after normalization to the β-actin loading control for each protein followed by normalization to the reference protein using Fiji/Image J.

### Flow cytometry

For detection of β1 integrin levels, cells were cultured as subconfluent monolayers and detached by trypsin-EDTA (2 mM), washed in PBS and stained with LIVE/DEAD stain (30 min, 4°C). 4T1 cells were stained with primary antibodies and matching IgG controls dissolved in FACS buffer (2% FBS/2mM EDTA in PBS; 1 hour, 4 °C), washed with FACS buffer, incubated with secondary antibody dissolved in FACS buffer (30 – 60 min, 4 °C), and analyzed by flow cytometry (BD FACS CANTO II; BD Biosciences). Quantification of the median fluorescence intensity (MFI) was calculated by linear subtraction of IgG control MFIs followed by total β1 integrin content and/or normalization to MFIs of DMSO, as indicated. All data analysis was performed using FlowJo (version 10.3.).

### Tumoroid invasion assay in 3D collagen matrices

Multicellular tumoroids were generated from subconfluent culture using the hanging-drop assay (20% methylcellulose; 500 cells/25 µL drop; overnight tumoroid assembly).^75^ Tumoroids were embedded in mid-density non-pepsinized rat-tail collagen type I (2.5 mg/ml, average pore size 4 µm^2^), low-density bovine collagen (1.7 mg/mL, 20 µm^2^), or high-density rat-tail collagen (4 mg/mL, 1 µm^2^).^12^ After collagen polymerization (37 °C, 10 – 20 min), 3D cultures were treated with DMOG (1 mM) or DMSO (0.1%, vehicle control) dissolved in respective cell culture media or placed in incubators with 21% normoxic or 0.2% hypoxic oxygen levels. Migration morphology and efficacy were monitored by brightfield time-lapse microscopy (Leica) or endpoint analysis after 72 hours (4T1) or 96 hours (UT-SCC38).

For cell counting of tumoroid 3D invasive cultures, tumoroid-collagen droplets were incubated with collagenase type I at 37 °C and cells were disaggregated by trypsin-EDTA (2mM).

### Pharmacological treatment of 3D tumoroid migration cultures

The molecular dependencies of cell migration modes and the migration efficacy of individual cells were tested in 3D collagen cultures after 48 hours (4T1) or 72 hours (UT-SCC38) in DMSO/DMOG or normoxia or hypoxia conditions. At these time points, single-cell migration was established. The following reagents and compounds were added to the culture media: β1 integrin–blocking antibody 4B4 (0.5-5 µg/mL for reducing speed; 10 µg/mL for probing morphology), β1 integrin–activating antibody TS2/16 (20 µg/mL, UT-SCC38), β1 integrin activating antibody 9EG7 (5 µg/mL, 4T1), calpain inhibitor PD150606 (50 or 100 µM), and etoposide (200 µM). Tumoroid cultures in 3D collagen were incubated (37°C) with compound and monitored by single time-point or, for up to 20 hours, time-lapse brightfield microscopy (Leica). All antibodies and inhibitors used in live-cell assays were purchased as azide-free compounds or dialyzed (Slide-A-Lyzer mini dialysis device, 20 MWCO).

### Transient transfection of cell monolayers and 3D tumoroid cultures

Cells in monolayer culture were incubated (20 hours, 37°C) with a mission-siRNA transfection mix (100 nM siCAPN2 or siControl, 128 μL MISSION siRNA transfection reagent, 1.2 mL serum-free medium, 6.8 mL complete medium, 200,000 cells, per 10cm^2^ dish) or with a plasmid DNA transfection mix (6 μg GFP-TalinL432G or pEGFP plasmid DNA, 15 µL Lipofectamine 2000, 500 µl OptiMEM, 9 mL complete medium, 75,000 cells, per 10 cm^2^ dish). The transfection medium was removed and transfected cells were used for further analysis.

For molecular intervention followed by 3D matrix culture, cell transfection was performed during 3D tumoroid aggregation. A mission-siRNA transfection mixture (100 nM of mission-siRNA siCAPN2 or siControl, 32 µL MISSION siRNA transfection reagent, 200ul serum-free medium) or a plasmid DNA transfection mixture (4 µg GFP-TalinL432G or pEGFP plasmid DNA, 10 µL Lipofectamine 2000, 75 µL OptiMEM) was added to the cell suspension (50,000 cells, suspended in aggregation medium containing 20 % methylcellulose) plated as hanging drops (500 cells per 25 µL per drop) for cell aggregation (18 hours, 37°C). Tumoroids were washed with medium and used for 3D collagen culture.

### Time-lapse microscopy

3D tumoroid cultures were placed and maintained (37°C) in wax-sealed migration cambers.^12^ Brightfield time-lapse microscopy was performed using DM IL LED microscopes using 20x/0.30 HI Plan objectives (Leica Microsystems), equipped with STC-405 cameras (Sentech). Image acquisition was obtained by the 16-Channel Recorder software (SVS-Vistek GmbH 2004) at 4 min intervals.

### Immunostaining and confocal microscopy

For immunofluorescence staining, 3D collagen cultures (100 µL) were fixed (2% PFA, 5 min, 37°C; followed by 4 % PFA, 10 min, 37°C), washed (PBS, 3×10 min), permeabilized and blocked (PBS/0.1 % BSA/10 % normal goat serum /0.3 % Triton X-100, 1 hour, RT), and incubated with primary antibody (in 0.1 % PBS/BSA/0.3 % Triton X-100, 18 hours, 4°C). Cultures were washed eight times (PBS, 1 hour each), incubated with DAPI and secondary antibody (18 hours, 4°C in PBS/0.1 % BSA/ 0.3% Triton X-100), washed three times (PBS, 10 min each), and imaged as whole-mount 3D samples. Confocal fluorescence and reflectance microscopy were performed by sequential single-channel confocal scanning using a CS APO 40x NA 1.15 objective (TCS SP5 Confocal Microscope, Leica Microsystems) and an inter-slice distance of 2 µm.

For detection of active β1 integrin levels in situ, 3D collagen cultures (100 μL) of 4T1 tumoroids were incubated with active β1 integrin antibody 9EG7 diluted in medium (15 min, 37°C), washed (warm PBS, 3×5 min each) and fixed in 2% PFA (5 min, 37°C) followed by 4 % PFA (10 min, 37°C). Samples were blocked (1 hour, blocking buffer: 0.1 % PBS/BSA/10% normal goat serum) and incubated with DAPI, phalloidin, primary and secondary antibody dissolved in blocking buffer (18 hours, 4°C). Structured illumination microscopy using a 20x/0.8 objective (Zeiss LSM880/AiryScan) was performed for detecting phalloidin, active β1 and total β1 integrin signals at an inter-slice distance of 1 µm.

### Calpain activity assays

For detection of calpain activity in subconfluent monolayers, cells were detached with trypsin-EDTA (2mM), washed (PBS), incubated in the dark with LIVE/DEAD dye (Thermo Fisher Scientific) together with CMAC (10 µM, 20 min, 37°C), washed (PBS), incubated with calpain inhibitor PD150606 (100 µM in medium, 2 hours, 37°C) or DMSO vehicle and fixed (0.5% PFA/PBS, 18 hours, 4°C). CMAC substrate cleavage was detected by flow cytometry (LSRII, BD) using 355nm laser excitation and detected through a 440/40 filter (excitation/emission peaks at 351/430 nm). As reference for baseline fluorescence, cell autofluorescence was used and subtracted from the MFI of cleaved CMAC intensity using FlowJo (version 10.3). Calpain activity in 3D tumoroid culture or in human tumor xenografts was detected using multiphoton microscopy (LaVision BioTec) using a 25x NA 1.05 water objective (Olympus). Live tumoroid-containing 3D collagen cultures were washed (PBS, 5 min), incubated with CMAC (20 µM, 30 min, 37°C), and fixed (2% PFA in PBS, 5 min, 37°C; followed by 4 % PFA, 10 min, 37°C). For calpain inhibition, 3D tumoroid cultures were incubated with calpain inhibitor PD150606 (100 µM, 37°C, 20 hours), and inhibitor was refreshed 2 hours before fixation. For shape analysis of CMAC-stained cells, 3D samples were incubated with AlexaFluor 546-conjugated phalloidin (1 hour, RT). Calpain activity in human tumor xenografts was detected in fresh-frozen cryosections (50 μm), which were thawed (5 min, 20°C), incubated with CMAC (100 µM, 45 min, 37°C), washed twice (PBS, 5 min each) and fixed (2% PFA, 5 min, 37°C followed by 4% PFA, 10 min, 37°C). Images were acquired as 3D stacks with 2 μm inter-slice distance using excitation at 730 nm (20 mW power under the objective) and simultaneous 2-channel emission of cleaved/uncleaved CMAC was obtained using customized 427/10 filter (Semrock) and dichroic filter 405 nm (Semrock), respectively. Further channels were AlexaFluor 546-conjugated phalloidin (excitation: 1090nm; detection: 620/50), second harmonic generation (excitation: 1090nm; detection: 525/50).

### Metabolism in 3D culture

Oxygen consumption rates (OCR) and extracellular acidification rates (ECAR) in 3D tumoroid invasion cultures were quantified using the Seahorse Bioscience XFe24 analyzer (Agilent). Individual 4T1 tumoroids were placed in collagen droplets (rat-tail collagen, 4 mg/mL) in XF24 V28 plates (Agilent) and treated with DMOG (1 mM) or solvent (DMSO) 48 hours prior to metabolic analysis. Thirty minutes before the measurements, the culture medium was replaced by XF Base medium (Agilent) supplemented with 2 mM L-glutamine, 1 mM pyruvate, 12 mM glucose (pH 7.4, 37°C), and OCR and ECAR were measured for 2 min periods with 3 min wait intervals. Three basal activity measurements were followed by treatment with oligomycin (2 µM), FCCP (1 µM), and a combination of Rotenone (2 µM) and Antimycin-a (2 µM) in triplicate measurements.

For detection of lactate release, tumoroid invasion cultures were maintained (48 hours) under normoxic and hypoxic conditions or treated with DMOG (1 mM) or control vehicle (DMSO). The supernatant was harvested, treated with sodium oxamate (100 µM, LDH inhibitor) to prevent lactate-to-pyruvate conversion, and secreted lactate was determined colorimetrically (Lactate kit K607; Biovision). In brief, a reaction mix containing the sample, enzyme mix and buffer was generated and incubated (20°C, 30 min, in darkness). Absorbance was measured at OD 570 nm and corrected by subtracting the background absorbance and the basic lactate-supplement from the medium and was normalized to the number of tumoroids.

### Live-cell microscopy of mitochondrial activity in 3D culture

To evaluate the mitochondrial membrane depolarization status, JC-1 dye (10 μg/mL) or solvent (DMSO) were added (for 60 min) to 3D invasion culture. For pharmacological uncoupling of mitochondrial oxidative phosphorylation, FCCP (200nM) or solvent (ethanol, 1:5,000) was added (40 min after start of JC-1 incubation, for 20 min total). Live tumoroid cultures were washed (HBSS, 3 x 5 min) and confocal microscopy (LSM880, Zeiss; 37°C) was performed in phenol-red free medium as 3D stacks (2 μm inter-slice distance) with the following settings: 488nm excitation, detection (Ch2GaAsP detector) at 511 to 567 nm (green) and 575 to 624nm (red).

### Image analysis and single cell cytometry of 3D culture

Brightfield, confocal, and multiphoton microscopy images were analyzed and processed using Fiji/ImageJ (version 1.51). Excluded from analysis were: dead cells, detected by cellular or nuclear fragmentation or condensation; mitotic cells, detected by cell swelling and/or rounding; and post-mitotic cells, including oppositely polarized cell doublets.

Protrusion types and number and the frequency of migrating single cells exhibiting a classified protrusion type were counted using the phalloidin channel from confocal maximum intensity z-projections (40x/1.15 objective). Migration morphology was classified from brightfield movies or still images (20x/0.30 HI Plan objective) acquired as z-series from non-overlapping tumoroid quadrants (4T1) or whole tumoroids (UT-SCC38). Scoring was performed by two independent operators in a non-blinded fashion and results of selected experiments were verified by double-blinded analysis. The total number of 4T1 subtypes per tumoroid was obtained from tumoroid quadrants and multiplied by 4. Migration paths and speed of single cells were digitized using Autozell (software version 080912; Center for Computing and Communication Technologies [TZI], University of Bremen, Bremen, Germany). The average migration speed was calculated by the length of the xy-path divided by the time. Only viable cells were included in the analysis. Cell elongation of detached cells was measured from stand-still images as the ratio of the maximum cell length over the maximum cell width.

The activation status of β1 integrin on individually migrating cells in 3D collagen culture was obtained by manually segmenting detached cells, creation of the summed intensity z-projection, and automated segmentation of the nucleus (DAPI channel) and F-actin (phalloidin channel). The merged DAPI and phalloidin channels were used for defining the cell area from which the raw integrated density of both active and total β1 integrin epitope was quantified.

Calpain activity in invading cells was quantified from maximum intensity z-projections. Single detached cells were segmented manually from the phalloidin channel using the shape tool. Cleaved and uncleaved CMAC substrate intensities were calculated as corrected total cell fluorescence (CTCF) with the formula: CTCF = Integrated Density – (Area of selected cell X Mean fluorescence of background readings). The signal ratio of cleaved/uncleaved CMAC was proportional to cleaved CMAC signal intensities only (427/10 emission), thus representing calpain activity by single channel detection. To identify calpain activity in migration subsets, CTCF per cell and cell elongation (maximum cell length/maximum cell width; phalloidin channel) were co-registered.

Mitochondrial membrane depolarization in invading single cells was obtained using JC-1. Summed intensity z-projections were created and the region of interest (single detached cells) were drawn manually using the shape tool on the brightfield channel. The mean intensity of the red and green JC-1 channels per cell were obtained to calculate the JC-1 ratio (red/green intensity).

### Computational modelling of cell elongation

A two-dimensional phase field model was used to simulate cells with different contractility and cell-substrate friction, which jointly regulate the cell shape, actin flow field, and traction force toward the substrate.^33^ In this model, cell contractility and cell-substrate friction lead to an elongated cell shape. The cell boundary was defined by a set of partial differential equations for the phase field ϕ which minimize its energy. For a single cell, its Hamiltonian (energy) can be written in form of equation (1):

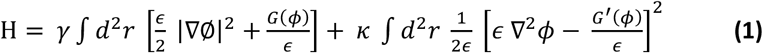

where the first term is related to the cell perimeter, and the second term represents the energy for the curvature integrated over the membrane. The phase field evolves with time, simulating the movement of the cell via coupling of the acto-myosin cytoskeleton and adhesion to the substrate to the phase field. This coupling involves active force generation through both actin polymerization and the activity of myosin motors. The detailed dynamics for the phase field systems underlying our simulations were taken from Shao et al.^33^

To regulate the intracellular contractility and friction between the cell and the substrate, three parameters were used: myosin contraction coefficient 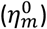, effective viscosity of actin flow (υ_0_), and gripping coefficient 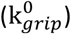. A larger 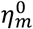 will lead to a larger contractility, and a larger 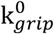 (or υ_0_) results in a larger friction. We assume a relationship between υ_0_ and 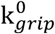 in the form of equation (2) to simplify the model,

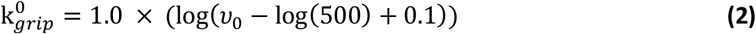

For each run, the simulation was initialized with different 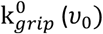 changes according to equation (2) and 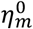. After initialization, the cell starts to move according to the dynamic equations. Cell shape was recorded at steady state. The elongation factor of cells with different 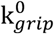 and 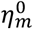 was calculated based on images as the ratio cell length/width vs. 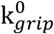 and 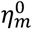 and represented as 3D landscape plot. Mathematical algorithms are unpublished but can be provided upon reasonable request.

### Orthotopic human HN-SCC xenograft model

Animal experiments were approved by the Animal Care and Use Committee (ACUC) of the University of Texas MD Anderson Cancer Center (00001148-RN00 and 00001522-RN00). Orthotopic human HN-SCC xenografts in nude mice were established as described.^76^ HN-31 and Detroit 562 cells were injected (5 x 10^4^ in 30 µL serum-free DMEM; Gibco) into the dorsal part of the tongue of 6- to 8-week-old male athymic nude mice (ENVIGO). Mice were euthanized by CO_2_ asphyxiation 30 days post-injection or when losing more than 20% of their pre-injection body weight. Tongues containing primary tumors were resected at the endpoint and cryopreserved.

### Tumor cell implantation *in vivo* and intravital multiphoton imaging

Intravital microscopy protocols were approved by the Animal Care and Use Committee (ACUC) of the University of Texas MD Anderson Cancer Center (00001002). A dorsal skin-fold chamber containing an optical imaging window was transplanted onto 8- to 10-week-old male athymic nude mice (MD Anderson Housing and Breeding Center in the Experimental Radiation Oncology Department, MD Anderson Cancer Center). Prior to injection, UT-SCC38 or UT-SCC58 cells expressing H2B-mCherry/Lifeact-GFP were cultured in the presence of DMOG (1 mM) with or without calpain inhibitor PD150606 (100 μM) for 48 hours. DMOG and PD150606 were refreshed after 20 hours and 2 hours before cell harvesting. Both weakly and firmly adhesive cells were harvested (PBS rinse, trypsin-EDTA (2mM) respectively), filtered through a FACS cell strainer cap (35 μm, Thermo Fisher) and injected as individual cells (1×10^4^ cells in PBS/mouse) intradermally into the optical imaging window 2 days after transplantation surgery. Four hours (UT-SCC58) and 20 hours (UT-SCC38) post-injection, intra-vital multiphoton microscopy was performed (3D z-stacks, 2 μm interslice distance, 3-5 regions, 1 animal/ cell line) with the following excitation and detection settings: Lifeact-GFP (920 nm, 20mW, BP 525/50); H2B-mCherry (1090 nm, 50mW, BP620/50); second harmonic generation (SHG; collagen) (1090 nm, 50mW, BD 525/50), third harmonic generation (1180 nm, 70mW, BP 387/15). Image analysis, 3D reconstruction and maximum intensity projections were performed using Fiji/ImageJ (version 1.51). Dying cells (nuclear fragmentation) and non-migrating cells (lacking actin-rich protrusions) were excluded from the analysis.

### Experimental lung metastasis

Animal experiments were approved by the Animal Care and Use Committee (ACUC) of the University of Texas MD Anderson Cancer Center (00001002). 2 days prior to tail-vein injection, cells were cultured under normoxic or hypoxic conditions in the presence of calpain inhibitor PD150606 (100 μM) or vehicle control (0.1% DMSO). PD150606 was refreshed after every 20 hours. Two hours after the last refreshment, weakly and firmly adhesive cells were harvested (PBS rinse, trypsin-EDTA [2mM] respectively), filtered through a FACS cell strainer cap (35 μm, Thermo Fisher) washed (PBS), and injected (30 G needle) as individual cells (500,000 cells in 100 μL/ mouse) into the tail vein of 8- to 10-week old female athymic nude mice (MD Anderson Housing and Breeding Center, Department of Experimental Radiation Oncology, MD Anderson Cancer Center). Fourteen days post-injection, mice were euthanized and subjected to whole-lung perfusion fixation (10 % neutral buffered formalin, 5 min). Lung macrometastases were manually counted, defined as whitish colonies on the lung surface. Metastasis by epithelial cells was immunohistochemically confirmed, by submerging the lungs in formalin (10%, 5-7 days) followed by sectioning (4 μm thickness; Research Histology Core Laboratory, MD Anderson Cancer Center). Antigen retrieval was performed in citrate buffer (Rodent Decloaker) and anti-cytokeratin 8 antibody (30 min incubation) was detected using rabbit-on-rodent HRP detection polymer (15 min incubation) visualized with diaminobenzidine, counterstained with hematoxylin and imaged with panomaric slide scanner (40X objective).

### Statistical analysis

Bar graphs represent the means and standard deviation and scatter diagrams the median and interquartile range (25%-75%) of individual cell data points from pooled experiments. Statistical analysis was performed using the two-tailed non-paired *t* test (two groups) or one- or two-way ANOVA (more than two groups). For non-parametric distributions determined by normality testing, Mann-Whitney U (two groups) or Kruskal-Wallis multiple comparison with Dunn’s correction (multiple groups) were used, unless otherwise indicated. Statistical analysis was performed using GraphPad Prism 8 software (GraphPad Software, Inc., San Diego, CA).

## Acknowledgements

We gratefully acknowledge Dr. Jeffrey N. Myers, Department of Head and Neck Surgery, MD Anderson Cancer Center for generous supply with samples from HN-31 and Detroit256 HNSCC tumor xenografts. We also gratefully thank Dr. Bettina Weigelin, Department of Genitourinary Medical Oncology, MD Anderson Cancer Center for help with intradermal injections of UT-SCC38 and UT-SCC58 cell lines. This work was supported by the European Research Council (617430-DEEPINSIGHT), NWO-Vici (918.11.626), the Cancer Genomics Center, and the MD Anderson Cancer Center Moon Shot program. V. te Boekhorst was supported by the Rosalie B. Hite Fellowship for Cancer Research. M. Mählen was supported by the Boehringer Ingelheim Fonds. Immunohistochemistry analyses were provided by the Tissue Biospecimen & Pathology Resource Core, supported by the NIH/NCI CCSG grant (5P30CA016672).

## Author Contributions

V.t.B designed and carried out the experiments, analyzed the data, supervised the work, and wrote the manuscript. L.J. performed immunoblotting, transfections, flow cytometry, tumoroid invasion culture and carried out morphology analysis. M. Mählen performed active β1 integrin and JC-1 experiments in 3D tumoroids. M. Meerlo and B.M.T.B. designed and executed metabolic analyses. F.C.D. generated β1 integrin and HIF-1α Western blots. Y.Y. and H.L. designed and carried out in silico analyses. G.D. performed viability, proliferation and colony formation analyses. P.F. designed the experiments, supervised the work, and wrote the manuscript. All authors read and corrected the manuscript.

## Conflict of Interest Statement

The authors declare no conflict of interest.

