## Supplementary information for "Calpain-2 regulates hypoxia/HIF-induced amoeboid reprogramming and metastasis"

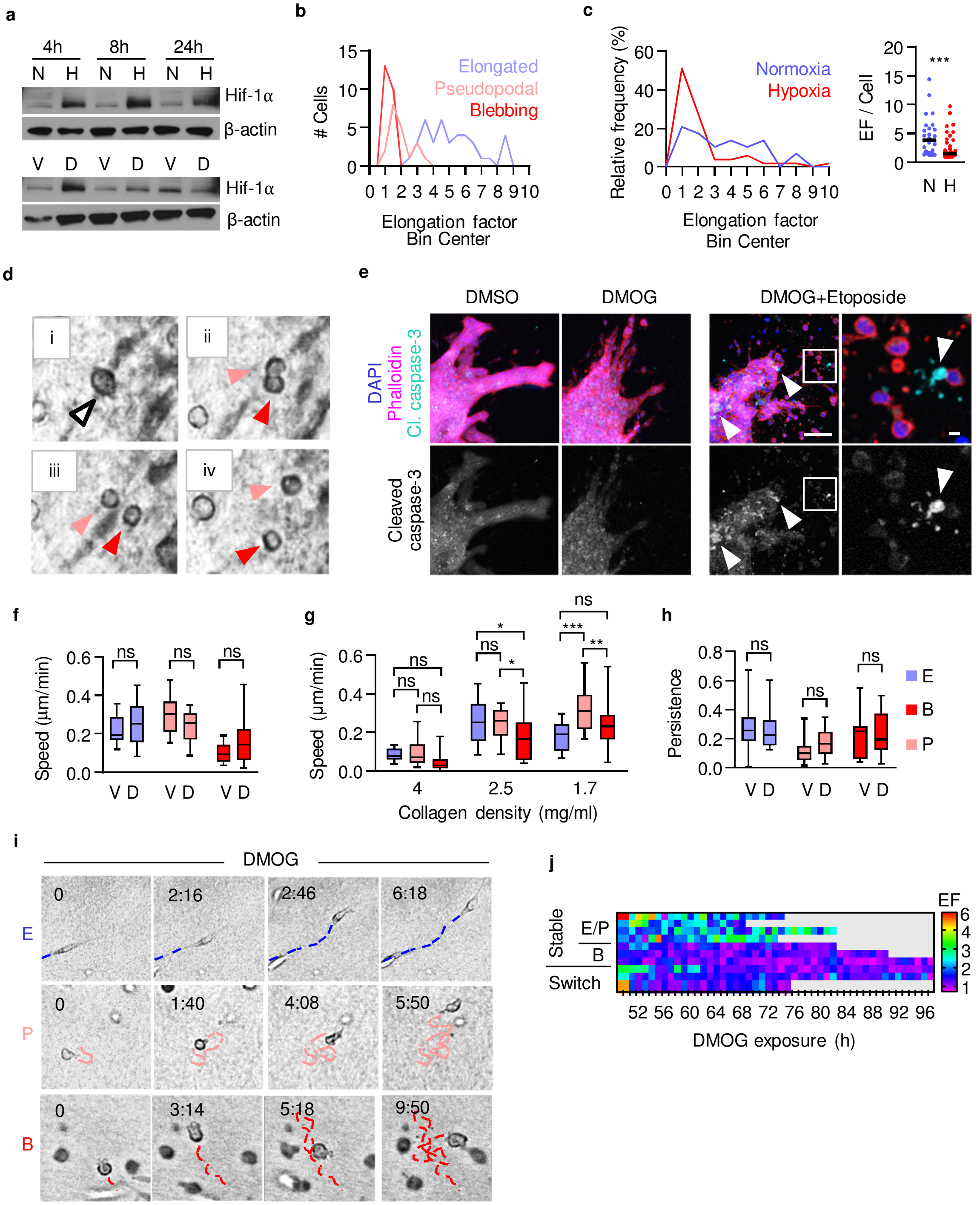


**Supplementary Figure S1. Morphological and functional mapping of migration modes induced by HIF stabilization. a,** Representative Western blot of HIF-1α protein levels in 4T1 cell monolayer culture. **b,** Elongation factor (EF) distribution of 4T1 single-cell migration subtypes (72 h; 86 cells, n=2). **c,** EF distribution of migrating 4T1 single-cells (left panel) and overall change of elongation (right panel) after HIF stabilization (72 h; 80 cells, n=2). Horizontal line, median. *** P=0.0002 (Mann-Whitney test, two-sided). **d,** Cell division of a blebbing-amoeboid migrating cell in collagen 72 h after DMOG treatment. Arrowheads, post-mitotic daughter cells. **e,** Confocal micrographs of cleaved-caspase 3 (apoptotic marker) in UT-SCC38 tumoroids in 3D collagen 96 h after HIF-stabilization. Etoposide treatment (200 µM) was used as positive control. Insets, blebbing amoeboid-migrating subtypes. Arrowheads, apoptotic cells positive for cleaved-caspase-3. Scale bars, 100 µm (overview), 10 µm (inset). **f,** Migration speed of individual 4T1 cells in intermediate-density collagen in the presence of DMOG or solvent. Box and whiskers show the median, 25/75 percentile and minimum/maximum (48-72 h; 37-60 cells, n=3). P-values: 0.58 (E); 0.15 (P); 0.27 (B) (unpaired t-test, two-sided). **g,** Migration velocities of 4T1 single-cells after HIF-stabilization with DMOG in collagen with low (1.7 mg/ml), intermediate (2.5 mg/ml) or high (4 mg/ml) density.

(48-72 h; 45 to 66 cells per density, n=3). Data are represented as in (**f**). *** $P=0.0001$, ** $P=0.009$, * $P<0.025$, ns, $P\geq 0.17$ (two-way ANOVA). **h,** Migration persistence. Conditions and data represented as in (**f**). P values: 0.87 (E); 0.45 (P); 0.99 (B) (unpaired t-test, two-sided). **i,** Migration paths and related shapes of DMOG-induced 4T1 cells in low-density collagen. Time (h:min). Scale bars, 10 µm. **j,** Elongation kinetics of individual UT-SCC38 cells during collagen invasion after HIF-stabilization. Rows represent one cell over time. Abbreviations: E, elongated-mesenchymal; P, pseudopodal-amoeboid; B, blebbing-amoeboid; N, normoxia; H, hypoxia; V, vehicle control (DMSO); D, DMOG.

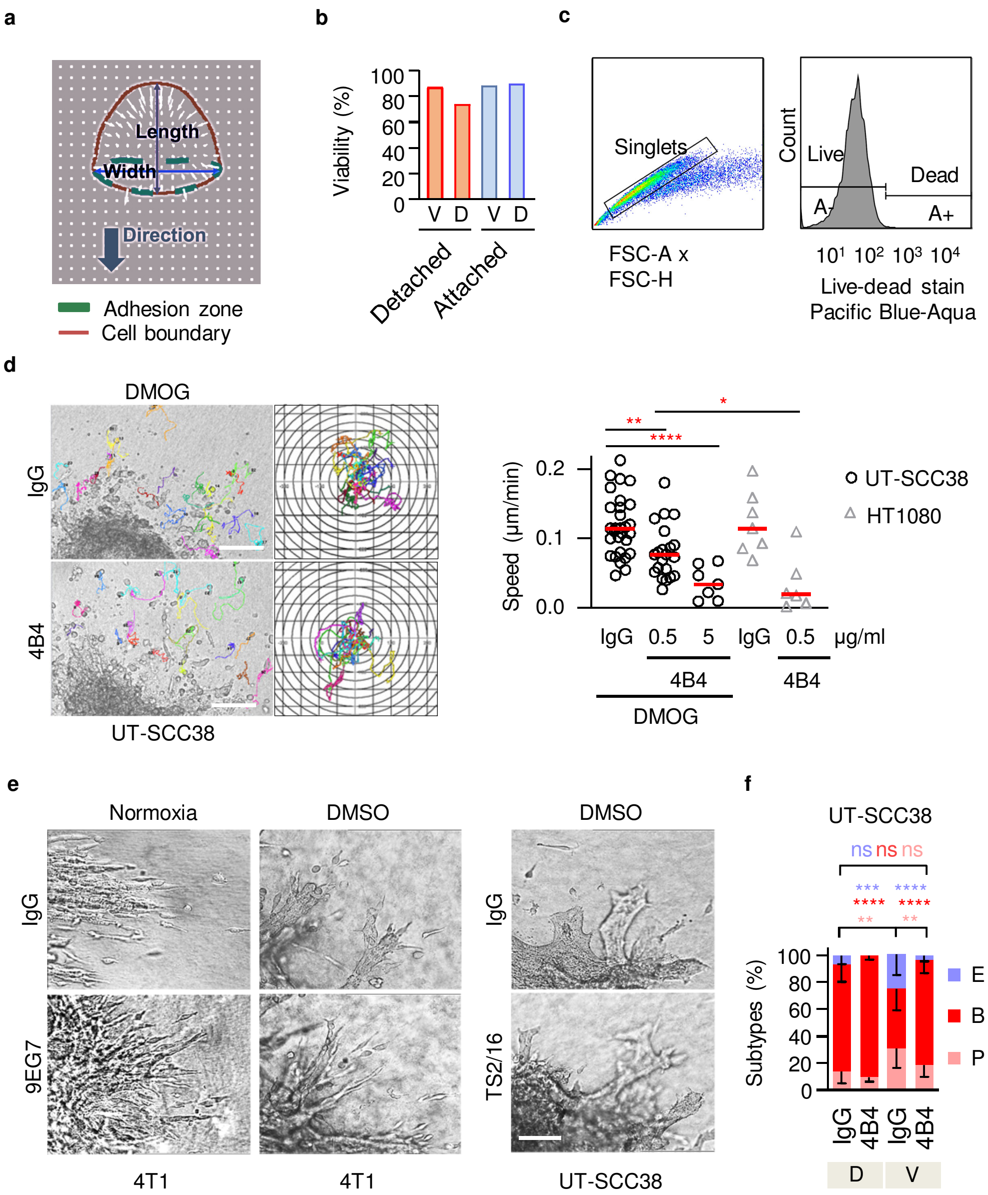


**Supplementary Figure S2. Cell viability, migration activity and integrin dependence of phenotypes after HIF stabilization. a,** In silico approach for shape simulation of migrating cells. Cell axes (length, width) used for elongation factor analysis. White arrows, velocity of actin flow. **b,** Cell viability of isolated 4T1 subsets 48 h after treatment in 2D culture, detected by Trypan blue exclusion. **c,** Flow cytometry gating strategy for selection of single viable cells (A- fraction) for receptor expression and activity analysis. **d,** Migration paths and trajectories (left panel) and speed (right panel) of individual UT-SCC38 cells after HIF-stabilization with or without β1 integrin blockade (mAb 4B4, 0.5 µg/ml; collagen 1.7mg/ml). For comparison, mesenchymal HT-1080 fibrosarcoma cells were exposed to the same treatment. Data show the median and individual cells (70 cells, n=2). **** P<0.0001, ** P=0.002, * P=0.01 (Mann-Whitney test, two-sided). **e,** Brightfield micrographs of migrating 4T1 cells in 3D collagen with or without β1 integrin activation during ongoing HIF-stabilization (4T1: 48-72 h, mAB 9EG7, 5 µg/ml; UT-SCC38: 72-96 h, mAb TS2/16, 20 µg/ml). **f,** Morphology-based scoring of single-cell migration subtypes after interference with β1 integrin activity (adhesion-perturbing mAb 4B4, 10 µg/ml). Data show the means ± s.d. from 49 tumoroids of two independent experiments. **** P<0.0001, *** P=0.0003, ** P<0.005 (two-way ANOVA). Abbreviations: E, elongated; P pseudopodal-amoeboid; B, blebbing-amoeboid; N, normoxia; H, hypoxia; V, vehicle (DMSO); D, DMOG. Scale bars, 100 µm.

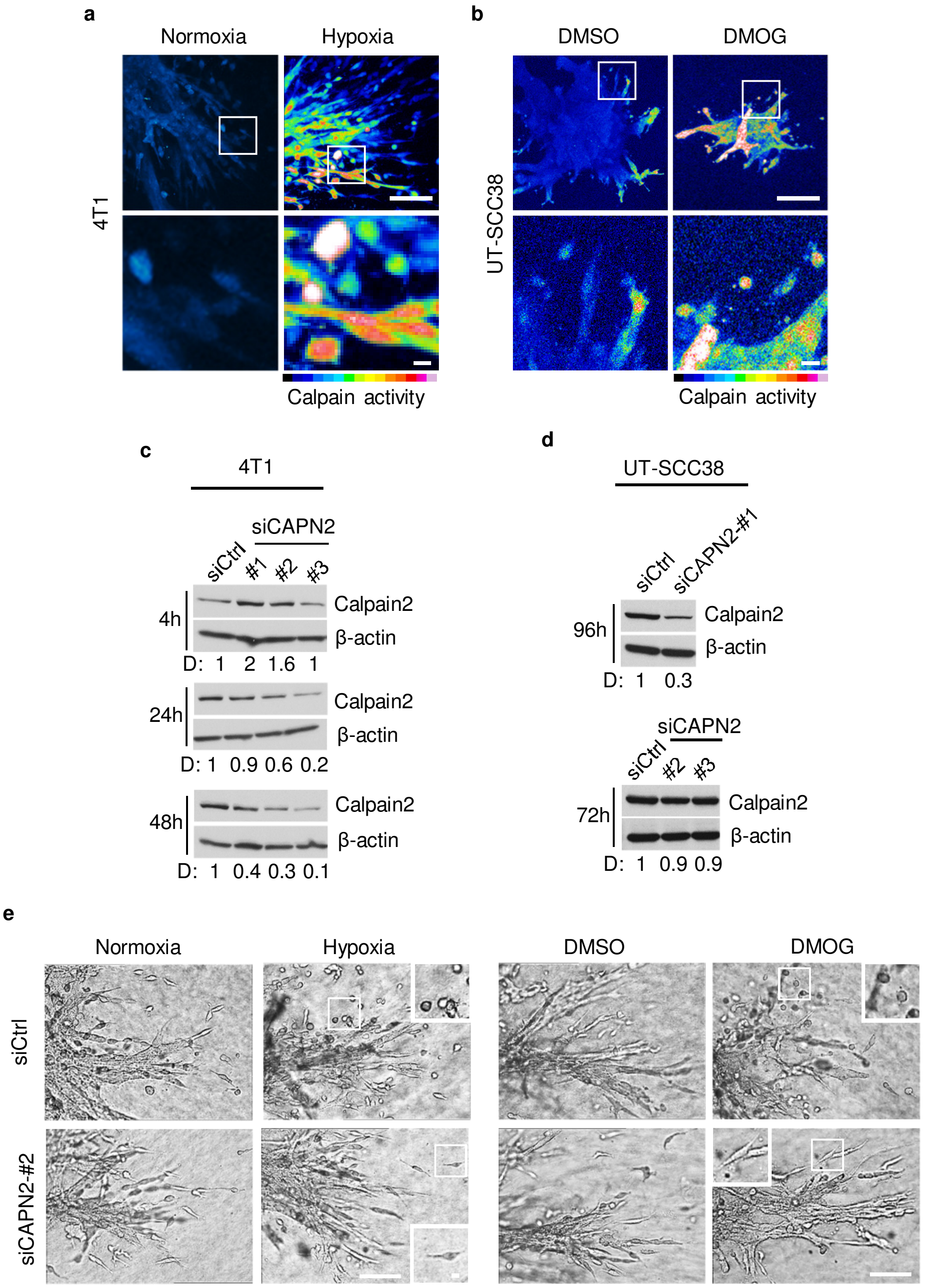


**Supplementary Figure S3. Calpain activity detection and RNA interference in 3D tumoroid invasion culture. a, b,** Calpain activity (cleaved CMAC intensity) of 4T1 tumoroids in 3D collagen after 72 h culture in normoxia and hypoxia (**a**) or in UT-SCC38 tumoroids 96 h after HIF-stabilization with DMOG (**b**). Representative multiphoton micrographs; insets, invasion zone. Scale bar, 100 µm (overview), 10 µm (insets). **c, d**, Calpain-2 protein expression after transfection with different siRNA targeting mouse calpain (CAPN2) (**c**) and human calpain-2 in UT-SCC38 cells (**d**). Representative Western blots and relative change compared to siCtrl determined by densitometry (D). For functional studies, mouse siCAPN2-#2 (4T1) and human siCAPN2-#1 (UT-SCC38) were pursued further. **e,** Migration morphologies of 4T1 tumoroids expressing siCtrl and siCAPN-2 72 h after transfection and simultaneous hypoxia/HIF-stabilization. Insets, single cell migration morphologies. Scale bars, 100 µm (overviews), 10 µm (insets).

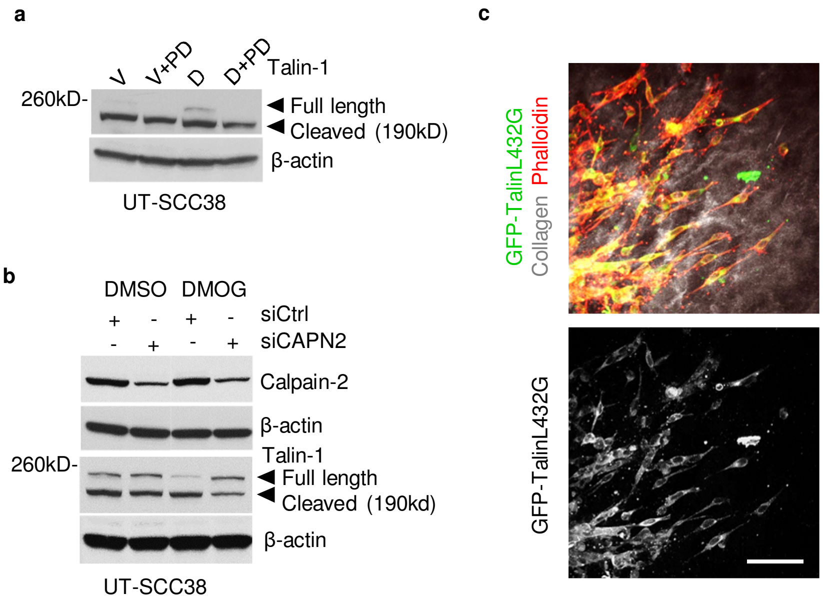


**Supplementary Figure S4. Biochemical and phenotypic analysis of tumor cells after expression of calpain-cleavage resistant talin. a,** Talin-1 cleavage after HIF-stabilization and siRNA-mediated (100nM) transient calpain-2 knockdown of UT-SCC38 cells. **b,** Transfection efficiency and transient expression of calpain-uncleavable GFP-TalinL432G. Representative multi-photon micrographs of 4T1 tumoroids in 3D collagen (SHG signal) 72h after transfection. Scale bar, 100 µm.

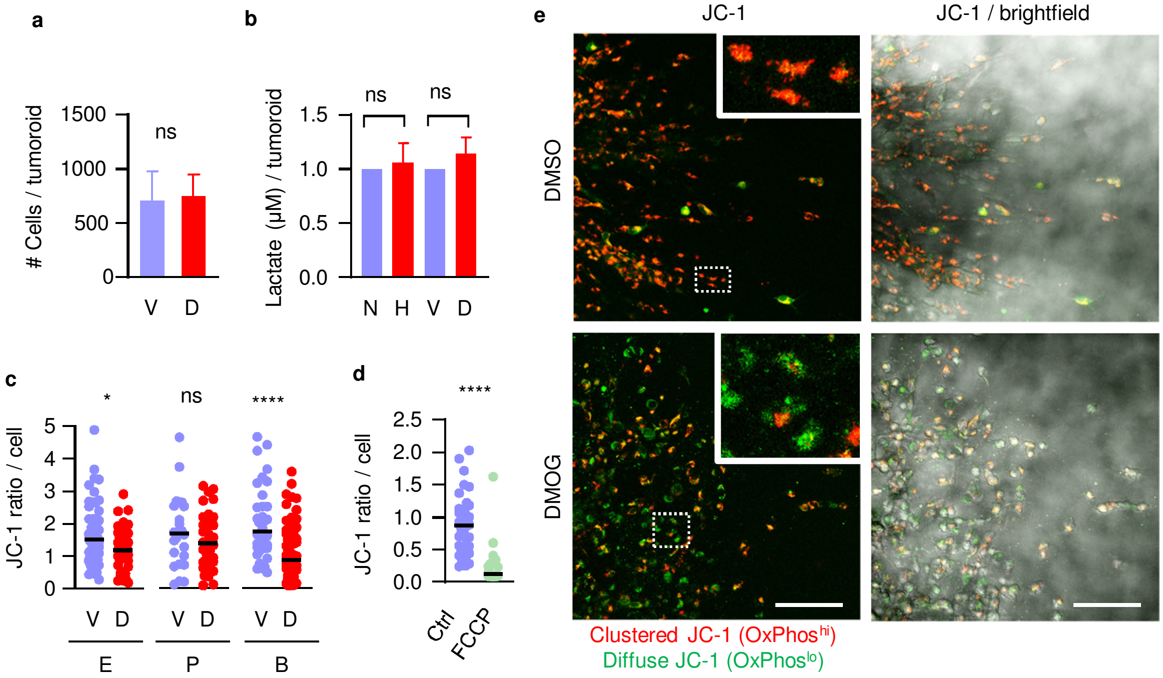


**Supplementary Figure S5. Standardization of cell number, lactate production and single-cell oxidative status in 3D tumoroid culture. a,** Cell number of individual 4T1 tumoroids isolated from 3D collagen droplets 48 h after DMSO or DMOG treatment used for Seahorse FX analysis. Data represent experimental means ± s.d. from 5-10 tumoroids per condition and experiment (n=3). ns, P=0.56 (Mann-Whitney test, two-sided). **b,** Average lactate levels in the supernatant of 4T1 tumoroids in collagen 48 h after HIF-stabilization. Data represent the experimental medians ± s.d. (n=3). Measurements from duplicate tumoroid-collagen droplets were averaged and normalized to the number of tumoroids. ns, P=0.99 (N, H) and P=0.31 (V, D) (Mann-Whitney test, two-sided). **c,** Subcellular quantification of JC-1 ratio in detached 4T1 single-cell subsets 72 h after HIF-stabilization with DMOG. JC-1 ratiometry was performed as described in in Fig. 5g. **** P<0.0001, * P=0.01, ns P=0.18 (unpaired t-test, two-sided). d, Subcellular quantification of JC-1 ratio in detached 4T1 single-cell subsets 72 h after normoxic culture treated with FCCP (200nM) or solvent (Ctrl). Data underlie the distribution curve in Fig. 5h. **** P<0.0001 (Mann-Whitney t-test, two-sided). **e,** Representative images (maximum projections; interslice distance, 2 µm) of 4T1 tumoroids in 3D collagen 72h after HIF-stabilization and incubation with fluorescent probe JC-1 60 min prior to microscopy. Scale bars, 100µm.

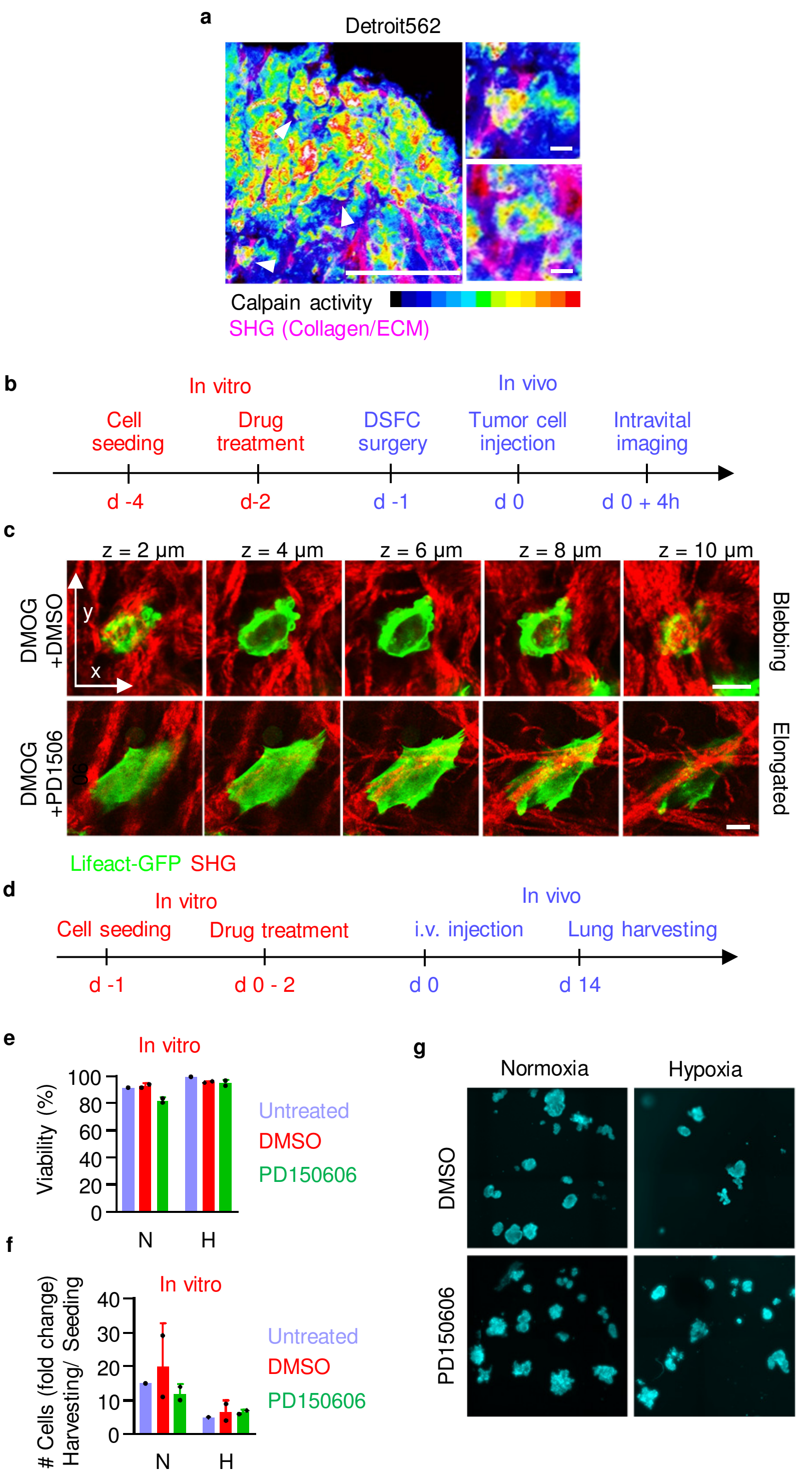


**Supplementary Figure S6. Calpain activity and function in vivo. a,** Calpain activity (cleaved CMAC) in ex vivo orthotopic human Detroit562 HN-SCC tumor xenograft tissue. Insets and arrowheads, invading single cells with round morphology positive for cleaved CMAC. Scale bars, 100 µm (overview), 10 µm (insets). **b,** Workflow for tumor cell implantation in vivo, including pre-treatment in vitro, injection and imaging of HN-SCC migration morphologies in the mouse dermis for monitoring of migration morphologies. No calpain inhibitor treatment was applied to mice. **c,** High-resolution z-projection of UT-SCC38 single-cells in the mouse dermis detected by multiphoton microscopy. Collagen interactions of a round cell with blebbing protrusions observed after DMOG pretreatment (top) and an elongated cell with podal protrusions after pre-treatment with DMOG combined with calpain inhibition (PD150606, 100 µM) (bottom). Scale bars, 10 µm. B, Blebbing amoeboid; E, elongated, mesenchymal. **d,** Workflow for the experimental lung metastasis assay, including pre-treatment in vitro, injection and harvesting of lungs at day 14. **e, f, g,** Viability (**e**), cell proliferation rates (**f**) and colony formation (**g**) of 4T1 cells after pre-treatment in 2D monolayer culture in vitro prior to tail-vein injection (workflow described in **d**). Data are mean ± s.d., with points representing independent experiments.

**Movie 1. Blebbing amoeboid single cell migration subtypes induced by DMOG.** Time-lapse brightfield microscopy of UT-SCC38 tumoroid invasion into 3D fibrillar collagen I in the presence of DMSO (solvent) or DMOG to pharmacologically stabilize HIF. Time (h : min). Field size 416 x 312 µm.

**Movie 2. Mesenchymal-to-blebbing amoeboid transition after detachment from tumoroids in the presence of HIF-stabilizing DMOG.** Time-lapse brightfield microscopy of 4T1 tumoroid invasion into 3D fibrillar collagen I. Arrowheads, initial and resulting subtype. Time (min). Field size 278 x 165 µm.

**Movie 3. Dependence of blebbing-amoeboid transition on reduced β1-integrin activity.** Time-lapse brightfield microscopy of 4T1 tumoroid invasion into 3D fibrillar collagen I in the presence of DMOG with β1-integrin activating mAb (clone 9EG7) or with IgG control. Time (h : min). Field size 416 x 312 µm.
